# Apical constriction induces tissue rupture in a proliferative epithelium

**DOI:** 10.1101/2022.03.02.482459

**Authors:** Mariana Osswald, André Barros-Carvalho, Ana M Carmo, Nicolas Loyer, Patricia C Gracio, Claudio Sunkel, Catarina C Homem, Jens Januschke, Eurico Morais-de-Sá

## Abstract

Apical-basal polarity is an essential epithelial trait controlled by the evolutionarily conserved PAR-aPKC polarity network. Deregulation of polarity proteins disrupts tissue organization during development and in disease, but the underlying mechanisms are unclear due to the broad implications of polarity loss. Here, we uncovered how *Drosophila* aPKC maintains epithelial architecture by directly observing tissue disorganization after fast optogenetic inactivation in living adult flies and ovaries cultured *ex vivo*. We show that fast aPKC perturbation in the proliferative follicular epithelium produces large epithelial gaps that result from increased apical constriction, rather than loss of apical-basal polarity. Accordingly, we could modulate the incidence of epithelial gaps by increasing and decreasing actomyosin-driven contractility. We traced the origin of epithelial gaps to tissue rupture next to dividing cells. Live imaging shows that aPKC perturbation rapidly induces apical constriction in non-mitotic cells, producing pulling forces that ultimately detach dividing and neighbouring cells. We further demonstrate that epithelial rupture requires a global increase of apical constriction, since it was prevented by the presence of non-constricting cells. Conversely, a global induction of apical tension through light-induced recruitment of RhoGEF2 to the apical side was sufficient to produce tissue rupture. Hence, our work reveals that the roles of aPKC in polarity and actomyosin regulation are separable and provides the first *in vivo* evidence that excessive tissue stress can break the epithelial barrier during proliferation.

## Introduction

Cell polarity is a defining feature of epithelial architecture and function. Apical-basal polarity ensures the asymmetric localization of the intercellular junctions that maintain the cohesion of epithelia and thereby preserve the epithelial barrier. Epithelial architecture is also regulated by the distribution of actomyosin-driven forces at the apical, basal and junctional level [1, 2]. It is thus not surprising that polarity disruption induces epithelial disorganization during animal development or disease [3-7]. This raises the importance of spatial cues provided by polarity regulators to build and support the three-dimensional structure of an organ. However, because polarity proteins are involved in many different processes that can ultimately affect tissue shape, how these proteins maintain epithelial architecture remains a critical longstanding question.

Interfering with polarity regulators in monolayered epithelia leads to different defects that disrupt epithelial integrity. These include the formation of multilayered tissue [4, 5, 8], ectopic lumens [5, 9, 10] and the appearance of gaps [7, 8, 11, 12]. Extensive characterization using loss or gain of function perturbations linked these defects to junctional disorganization, misoriented cell division, defective control of proliferation or misdiferentiation. However, direct observation of how an epithelium becomes disorganized upon disruption of polarity regulators is still missing, which prevents a clear understanding of how defects arise.

Atypical Protein Kinase C (aPKC) is part of the apical PAR complex (Cdc42-Par6-aPKC) and is a central regulator of animal cell polarity[13, 14]. It generates apical-basal asymmetry through phosphorylation of a number of polarity proteins, including Baz/Par3, Lgl, Par-1, Yurt and Crb. Their phosphorylation regulates local cortical binding through the modulation of multivalent protein interactions, homo-oligomerization [15-20], or simply by reducing electrostatic interactions with plasma membrane phospholipids [21, 22]. Apical-basal polarization ultimately positions belt-like adherens junctions (AJ) at the apical-lateral border where they mechanically link neighboring cells.

In addition to its well studied role in polarity regulation, aPKC regulates cell fate, epithelial-to-mesenchymal transition, cell cycle length, cell division orientation, actomyosin contractility and microtubule dynamics [9, 23-32]. In fact, some aPKC targets are not polarity proteins. Phosphorylation of Rho-associated coiled-coil-containing kinase (ROCK) represses the localization of this myosin activator to apical junctions, and thereby inhibits apical constriction in mammalian cells [26, 33, 34]. Moreover, aPKC acts as a negative regulator of apicomedial actomyosin networks to regulate pulsatile apical constriction in the *Drosophila* amnioserosa [35, 36]. aPKC function is also linked with actomyosin reorganization during cell division in fly tissues, which is consistent with its mitotic redistribution along the lateral cortex in mouse and sea anemone blastomeres [37-41]. Thus, aPKC may ensure epithelial architecture through different functional outputs, and assessing the contribution of actomyosin regulation and polarity maintenance demands separation of these aPKC functions.

Here, we combined optogenetic with chemical-genetic approaches to finetune aPKC inactivation with high-temporal control. This allowed us to disentangle the functions of aPKC in the regulation of actomyosin contractility and polarity. The monolayered follicular epithelium that encloses the *Drosophila* germline is a powerful system to explore the mechanisms that regulate epithelial architecture *in vivo*. Acute perturbation allowed us to capture that epithelial gaps form in the proliferative stages, where they arise by tissue rupture next to dividing follicle cells. We show this phenotype stems from the role of aPKC as an inhibitor of apical actomyosin networks in non-mitotic cells. These become hypercontractile after aPKC downregulation and pull on dividing cells until detachment occurs. Our work reveals the importance of keeping apical contractility in check during proliferation-mediated growth to maintain epithelial integrity.

## Results

### Optogenetic clustering inactivates aPKC

To explore how apical polarity maintains epithelial architecture, we developed a new approach to inactivate aPKC with high temporal control in the *Drosophila* follicular epithelium with an optogenetic clustering tool – light activated reversible inhibition by assembled trap (LARIAT) [42]. When exposed to blue light, LARIAT components – CRY2 fused to a GFP nanobody (V_H_H) and CIBN fused to a multimerization domain – interact with each other and cluster. To target and sequester aPKC, we used flies with endogenously GFP-tagged aPKC [43] and co-expressed GAL4-driven UAS-LARIAT (UAS-V_H_H::CRY2-P2A-CIBN::MP) [44] specifically in the follicular epithelium (Figure 1A).

**Figure 1.**
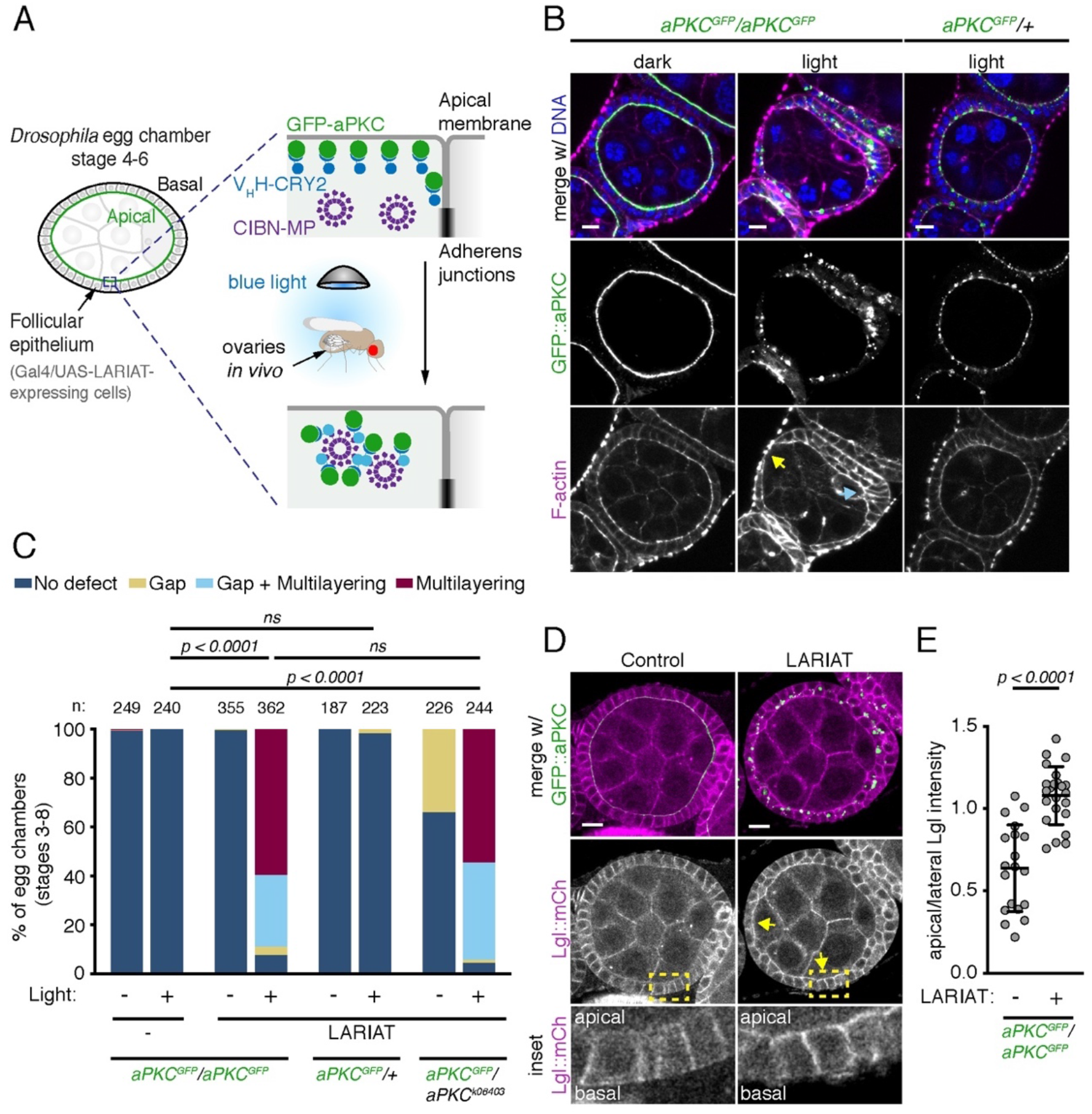
Optogenetic clustering inactivates aPKC and disrupts tissue architecture *in vivo*. **(A)** Schematic representation of optogenetic aPKC inactivation strategy using LARIAT (V_H_H::CRY2 and CIBN::MP). GFP::aPKC is targeted by CRY2 fused with a GFP nanobody (V_H_H). Exposing flies to blue light triggers CRY2 binding to CIBN fused with a multimerization domain (MP) and clusters GFP::aPKC. **(B)** Living flies were exposed to blue light for 48 hours to cluster GFP::aPKC or kept in the dark (control) before egg chambers were stained for F-actin and DNA. Control flies were kept in the dark. Flies were either homozygous or heterozygous for endogenously-tagged GFP::aPKC. Arrows point to epithelial gap (yellow) and multilayering (cyan). **(C)** Frequency of epithelial defects in egg chambers (stages 3 to 8) from flies with the indicated combinations of wild-type, GFP::aPKC or *apkc*^*k06403*^ null allele after 24 hours blue light exposure (n = number of egg chambers). LARIAT was expressed in the follicular epithelium when indicated. Control flies were kept in the dark. Fisher’s exact test compared the incidence of defects between different samples (ns, not significant). **(D)** Representative midsagittal images of control and LARIAT egg chambers from flies expressing GFP::aPKC and Lgl::mCherry exposed to blue light for 24 hours. Arrows point to apical Lgl::mCherry. Yellow boxes define region in insets. **(E)** Ratio of apical/lateral mean pixel intensity of Lgl::mCherry in control (n = 684 cells, 19 egg chambers) and LARIAT (n = 447 cells, 23 egg chambers). Graphs show mean ± SD, grey points represent average for individual egg chambers (t-test). Scale bars: 10 µm. See also Figure S1.

Expression of the UAS-LARIAT system in homozygous GFP::aPKC flies that remained in the dark did not interfere with aPKC localization or protein levels, and did not produce defects in epithelial organization (Figures 1B and S1A). This shows that GFP::aPKC is fully functional in the presence of the LARIAT components. We then exposed female flies to blue light continuously for at least 24h to test if optogenetic clustering reproduced the aPKC loss of function phenotypes described for the follicular epithelium [45, 46]. GFP::aPKC clustered in puncta, and led to the anticipated defects in epithelial architecture, namely epithelial gaps and multilayering (Figures 1B and 1C). A similar frequency of tissue defects was also visible after clustering heterozygous GFP::aPKC in the presence of an *apkc* mutant allele, but not in presence of the untagged wild-type allele (Figures 1B and 1C), which suggests that clustering inactivates GFP::aPKC. Furthermore, as predicted for aPKC inactivation, its substrate Lgl mislocalized to the apical domain upon aPKC optogenetic clustering in follicle cells (Figures 1D and 1E). Taken together, these results show that illuminating flies is sufficient to trigger CRY2-CIBN heterodimerization and perturb aPKC with optogenetics *in vivo* in adult flies prior to dissection. We further evaluated the impact of optogenetic aPKC clustering in the assymetric distribution of Miranda during *Drosophila* neural stem cell division, where it is a relevant aPKC substrate [47]. aPKC clustering prevented Miranda’s release from the apical domain of dividing larval neuroblasts (Figures S1B and S1C). Thus, LARIAT-mediated clustering is applicable to study aPKC in distinct contexts of cell polarity.

### Optogenetic aPKC inactivation leads to fast tissue disorganization

We took advantage of optogenetic perturbation *in vivo* to monitor the progression of tissue disorganization in flies exposed to light for specific periods of time (Figures 2A and 2B). We narrowed the analysis to stages 4 to 6 of egg chamber development, so as to determine the impact of aPKC perturbation in epithelial architecture prior to major morphogenetic changes during egg chamber development. Multilayering was the most prevalent phenotype in fixed tissue from flies exposed to light for longer periods of time, whereas gaps were the most frequent defect upon 2 hours of GFP::aPKC clustering *in vivo* (2h: ∼50% egg chambers with gaps and ∼30% with multilayering; 4h: ∼25% gaps and ∼85% multilayering; Figures 2A and 2B). The two phenotypes were not mutually exclusive (Figure 1C), and were commonly observed in different positions of the egg chamber (Figure 2C). Epithelial gaps appeared almost exclusively at the dorsal/ventral region, whereas multilayering was largely restricted to the egg chamber poles, which suggests a distict basis for the two phenotypes.

**Figure 2.**
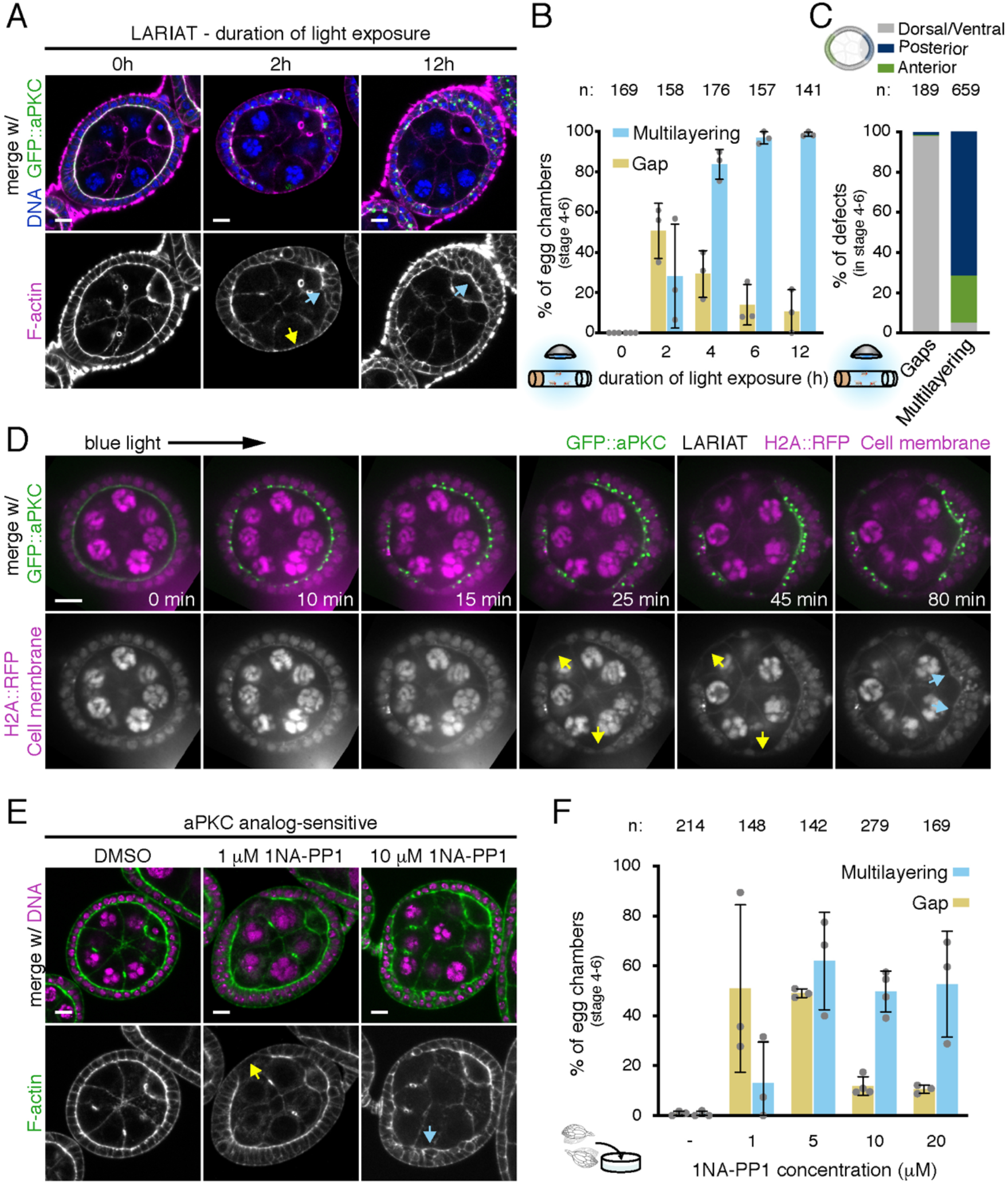
aPKC inactivation leads to fast tissue disorganization in a proliferative epithelium. **(A-C)** Representative midsagittal images and quantification of epithelial defects in GFP::aPKC homozygous egg chambers in proliferative stages (4-6) expressing LARIAT and stained for F-actin and DNA. Flies were exposed to light for the indicated time before ovary fixation. **(B)** Frequency (mean ± SD) of epithelial gaps (yellow arrow in A) and multilayering (cyan arrows in A). **(C)** The data from (B) was re-analyzed to show the spatial distribution of all defects, n = total amount of gaps or multilayering events. **(D**,**E)** Time-lapse midsagittal images of an egg chamber expressing LARIAT (in the follicular epithelium), GFP::aPKC and H2A::RFP show epithelial gaps (yellow arrows) and multilayering (cyan arrows). Imaging with 488nm laser triggered LARIAT clustering from min 0 onwards. **(E**,**F)** Midsagittal images and epithelial defect quantification in proliferative *aPKC*^*as4*^ egg chambers treated *ex vivo* for 2 hours with the indicated concentrations of 1NA-PP1 to inactivate aPKC and stained for F-actin and DNA. **(F)** Graph shows frequency (mean ± SD) of epithelial gaps (yellow arrow) and multilayering (cyan arrow). Grey data points (graphs in (B,F)) represent independent experiments; n = number of egg chambers. Scale bars: 10 µm.

Intriguingly, gap frequency declined with increasing duration of light exposure (Figures 2A and 2B), suggesting that epithelial gaps appear specifically during the initial phase of aPKC clustering and before the formation of multilayered tissue. We confirmed these results by live imaging using fluorescent markers of the nucleus (H2A::RFP) and plasma membrane in egg chambers cultured *ex vivo*. GFP::aPKC formed large clusters at the apical domain within minutes of exposure to blue light (488 nm; Figure 2D; Movie S1). Epithelial gaps formed within 30 minutes of light exposure and earlier than tissue multilayering (Figure 2D; Movie S1).

aPKC is likely only partially inactive during the initial period of clustering due to the time necessary to completely cluster and mislocalize aPKC. Prolonged aPKC clustering could futher inactivate aPKC and favor multilayering. To test if the predominance of distinct defects was associated with the level of aPKC inactivation, we treated ovaries mutant for an *aPKC* ATP-analogue sensitive allele (*aPKC*^*as4*^) [48] for 2h with a range of 1NA-PP1 inhibitor concentrations. Treatment with 1μM of 1NA-PP1, which *in vitro* reduces aPKC activity to ∼15% [48], led predominantly to epithelial gaps (∼50% of egg chambers with gaps vs 10% with multilayering), whereas increasing drug concentrations led predominantly to multilayering (Figures 2E and 2F). Thus, epithelial gaps are associated with partial aPKC inhibition, whereas multilayering arises after strong loss of aPKC function. Moreover, epithelial gaps are the earliest defect in epithelial architecture after aPKC inactivation.

### aPKC antagonizes apical constriction in multiple *Drosophila tissues*

To identify the primary cellular effect underlying epithelial gaps, we clustered aPKC and analyzed the immediate impact on polarity, adhesion and the actomyosin cytoskeleton. In contrast to long-term clustering (Figure 1D), apical-basal polarity was not affected before gap formation, as both Lgl::mCherry and E-cad::mKate2 remained enriched at the lateral membrane and apical junctions, respectively (Figures 3A and S2A). However, the fluorescence of Sqh::mKate2, a tagged version of non-muscle myosin II regulatory light chain (MyoII-RLC), increased rapidly at the apical side of the epithelium within minutes of light-exposure and prior to the formation of gaps (Figures 3B, 3C; Movie S2). Furthermore, this apical myosin increase was accompanied by an increase in circularity of the apical surface of the epithelium (Figures 3B and 3D). This tissue deformation could result from alterations in the apical area of individual cells. Accordingly, live imaging of mosaic epithelia with clonal expression of UAS-LARIAT showed that optogenetic aPKC clustering induced constriction of the apical area of LARIAT-expressing cells (Figures 3E and 3F; Movie S3). Thus, upregulation of apical contractility is the earliest effect upon optogenetic aPKC inactivation.

**Figure 3.**
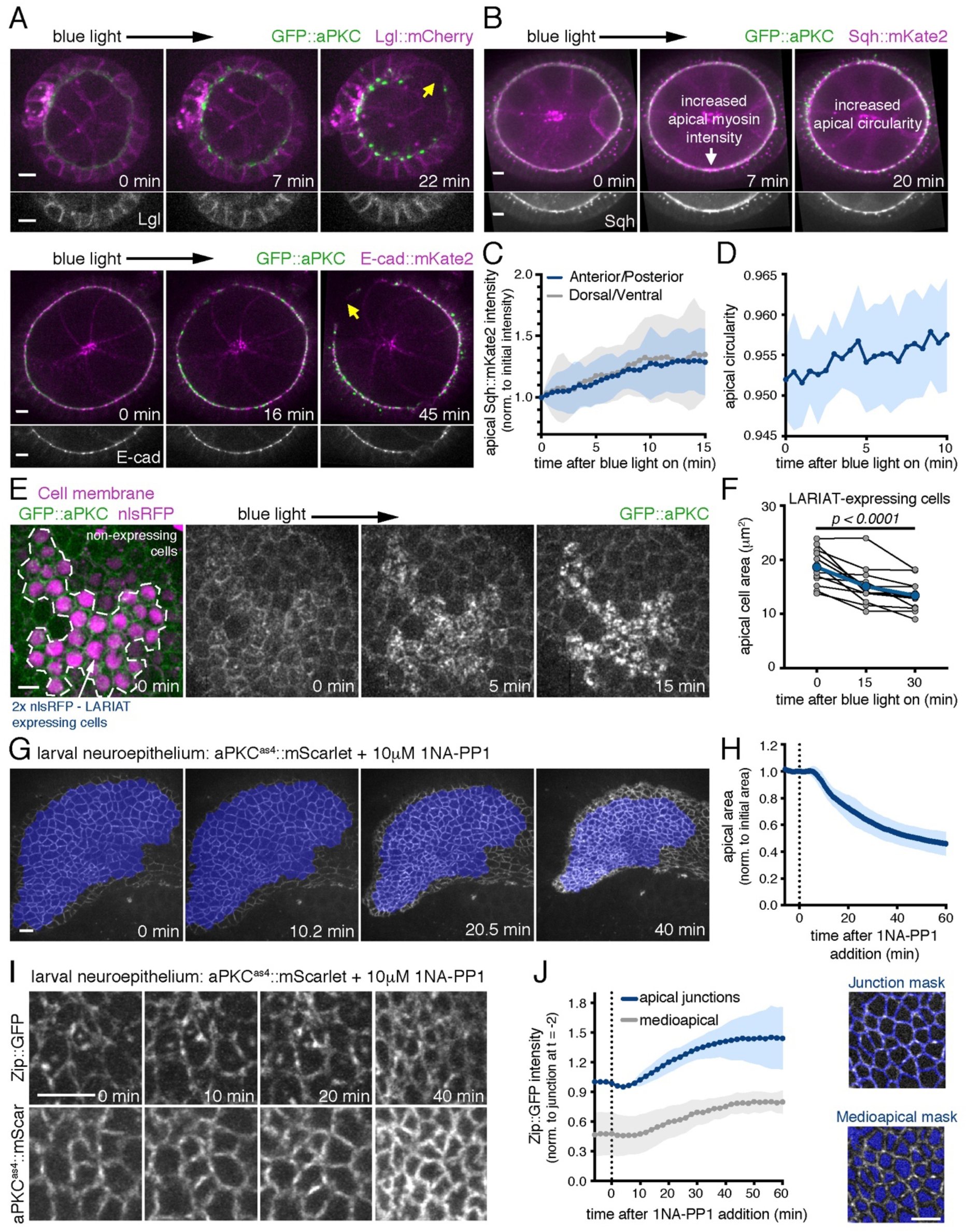
aPKC antagonizes apical constriction in *Drosophila* tissues. **(A, B)** Time-lapse midsagittal images of egg chambers expressing LARIAT, GFP::aPKC and either **(A top)** Lgl-mCherry, **(A bottom)** E-cad::mKate2 or **(B)** Sqh::mKate2. Imaging with 488 nm laser triggered aPKC clustering from min 0 onwards. Yellow arrows in (A) indicate epithelial gaps. **(C)** Sqh accumulation at the apical surface after aPKC clustering (mean ± SD) measured at the anterior/posterior (AP) and dorsal/ventral (DV) regions, corrected for cytoplasm intensity and normalized to its initial value (n ≥ 96 AP and ≥ 96 DV cells, 12 egg chambers). **(D)** Egg chamber circularity (mean ± SD) was measured at the apical surface of the follicular epithelium (n = 10 egg chambers). **(E)** Time-lapse images of GFP::aPKC follicular epithelium cells with mosaic LARIAT expression (marked by 2xnls::RFP in magenta). Imaging with 488 nm laser triggered aPKC clustering and apical domain constriction in LARIAT-expressing cells. **(F)** Average apical cell area (mean ± SD) within LARIAT expressing clones before and after aPKC clustering. Points represent average for individual clones (n = 161 cells, 12 clones, p<0001, ANOVA for paired samples). **(G)** Contraction of *aPKC*^*as4*^*::mScarlet* larval brain neuroepithelium following the addition of 10 µM 1NA-PP1 at min 0. **(H)** Graph shows apical area (mean ± SD, normalized to its initial value) of *aPKC*^*as4*^*::mScarlet* larval brain neuroepithelial tissue after addition of 10 µM 1NA-PP1 (n = 10 neuroepithelia with an average of 157 cells, 5 independent experiments). **(I)** Close-up of the neuroepithelium of an *aPKC*^*as4*^*::mScarlet* larvae expressing non-muscle myosin II heavy chain (Zip::YFP) following addition of 10 µM 1NA-PP1. **(J)** Zip::YFP intensity (mean ± SD) at the apical junctions and medioapical region (quantification masks shown) was normalized to junction intensity at -2 min (n = 4 neuroepithelia with ≥ 108 cells). Scale bars: 5 µm. See also Figure S2.

We further tested if the apical myosin increase was the first consequence of aPKC inactivation using chemical-genetics (Figures S2B and S2C). Live imaging showed that treatment of *aPKC*^*as4*^ egg chambers with 1μM 1NA-PP1 led to a quick increase of apical Sqh::mKate2, which persisted for 50 min in regions of the follicular epithelium without gaps. A higher inhibitor concentration (10 μM 1NA-PP1) also increased apical Sqh::mKate2 initially. However, this effect was transient, possibly due to the quicker loss of apical-basal polarity, which was reported to occur around 20 minutes after addition of 10 μM 1NA-PP1 in the follicular epithelium [48]. Perturbation of aPKC activity with high-temporal control therefore demonstrates that aPKC regulates apical contractility prior to polarity loss.

To determine if downregulation of apical myosin is a general function of aPKC, we analyzed the effect of aPKC inhibition on the neuroepithelium of the developing fly brain. Previous genetic perturbation in neuropithelial cells indicated that aPKC could support apical contractility by maintaining the polarized apical localization of myosin [28]. Therefore, we hypothesised that fast inactivation would also be necessary to separate the roles of aPKC in contractility and polarity in this tissue. We performed live imaging of *aPKC*^*as4*^::mScarlet brains to follow the initial impact of aPKC inhibition on apical shape and myosin accumulation. These experiments showed a dramatic constriction of the neuroepithelium within 10 minutes of inhibitor addition (Figures 3G and 3H; Movie S4). Apical constriction was associated with an increase in junctional and medioapical myosin II intensity (Figures 3I and 3J). Hence, aPKC downregulates apical constriction to control the shape of distinct epithelial tissues.

### Epithelial gaps result from increased apical contractility

To determine if increased apical actomyosin contractility is necessary to generate epithelial gaps in the follicular epithelium, we first disrupted the actin cytoskeleton with Latrunculin A (Lat A). Time-lapse imaging with E-cad::mKate2 to measure the apical area at the AJ level shows that treatment with LatA before light exposure blocks constriction during optogenetic clustering of aPKC (Figures 4A and 4B). Moreover, disruption of the actin cytoskeleton prior to aPKC clustering prevented the formation of epithelial gaps in tissue exposed *ex vivo* for 2 hours to light (Figures 4C and 4D). To further test whether increased MyoII activity promotes gap formation, we modulated actomyosin contractility and evaluated the presence of epithelial gaps after *in vivo* aPKC clustering for 2 hours. Overexpressed unphosphorylatable (SqhAA) or phosphomimetic (SqhEE) versions of myosin-RLC respectively reduce and increase contractility when they form bipolar filaments with wild-type myosin-RLC [49-51]. Upon optogenetic aPKC inactivation, SqhAA overexpression restored epithelial integrity, whereas SqhEE overexpression increased epithelial gap frequency (Figure 4E). Together, these results indicate that actomyosin-dependent cell contractility promotes and is necessary for the formation of epithelial gaps after aPKC inactivation.

**Figure 4.**
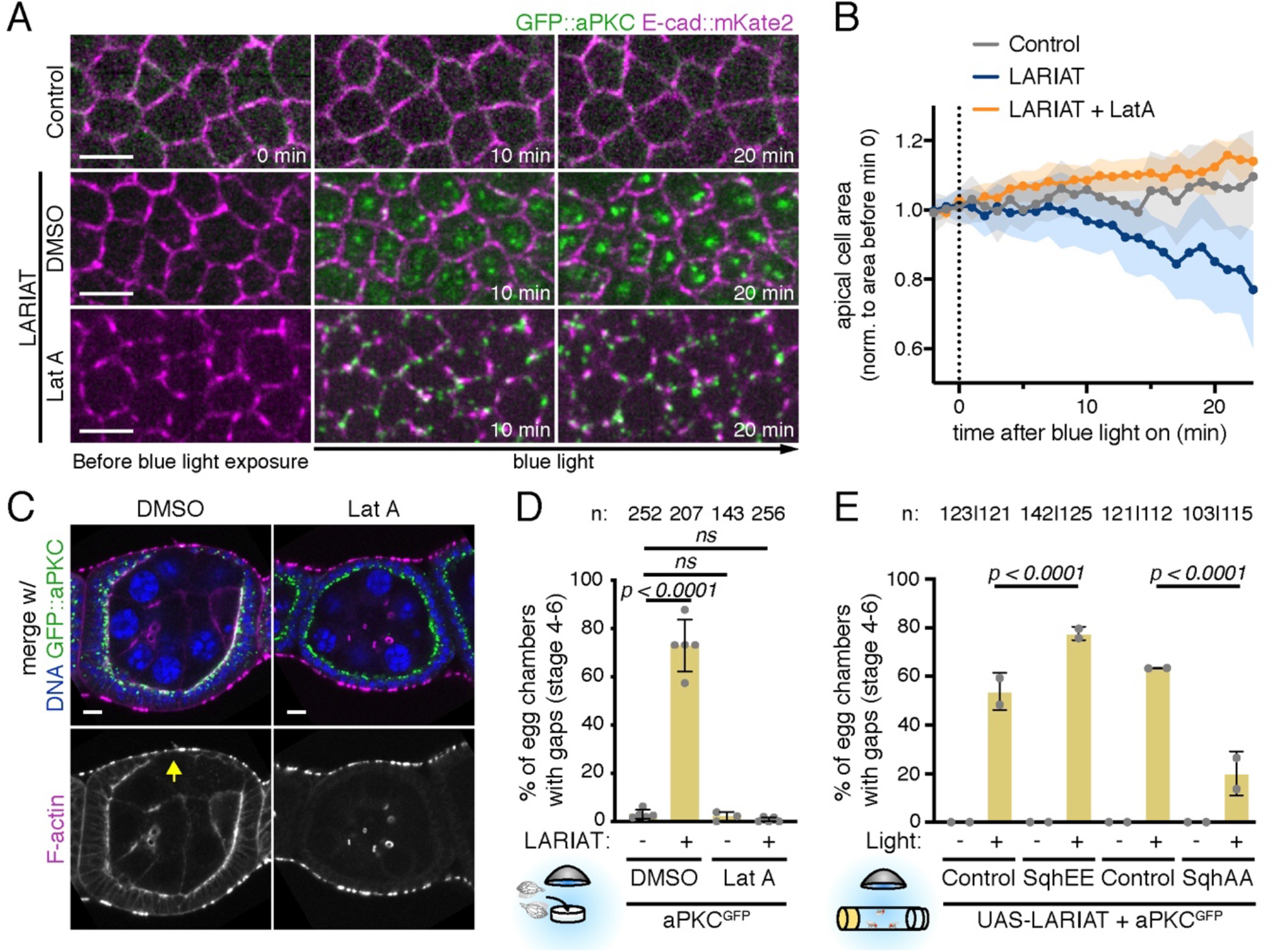
Increased apical contractility underlies gap formation after aPKC optogenetic clustering. **(A)** Live imaging of GFP::aPKC in control or LARIAT egg chambers co-expressing E-cad::mKate2 (apical view) and treated or not with Lat A prior to aPKC clustering (bottom). Imaging with the 488 nm laser triggered aPKC clustering from min 0 onwards. **(B)** Apical surface area (mean ± SD) measured at the junction level and normalized to the mean value before clustering (n ≥ 7 egg chambers per condition). **(C**,**D)** Representative midsagittal images and quantification of epithelial gaps in proliferative follicular epithelium expressing GFP::aPKC (green) and LARIAT and stained for F-actin and DNA. Ovaries were exposed to blue light *ex vivo* before fixation. Actin disruption by Lat A treatment restores epithelial integrity. **(D)** Frequency (mean ± SD) of epithelial gaps (yellow arrow in C) scored in the presence (+) or absence of LARIAT (-) in egg chambers treated with DMSO (control) or Lat A. **(E)** Epithelial gap frequency (mean ± SD) was scored upon overexpression of mCherry (Control), Sqh^E20E21^ (SqhEE) or Sqh^A20A21^ (SqhAA) in the follicular epithelium of GFP::aPKC LARIAT flies. Flies were exposed to blue light (+) or kept in the dark (-) for 2 hours prior to fixation; grey data points represent independent experiments; n = number of egg chambers scored; Fisher’s exact test (ns, not significant).

### Epithelial gaps form by tissue rupture next to dividing cells

During live imaging of Sqh::mKate2 in midsagittal egg chamber sections, we noticed that epithelial gaps frequently formed next to dividing cells (Figure S3A; Movie S5). To test if epithelial gaps appear specifically in proliferative tissue, we analysed egg chambers in stage 8 of oogenesis to ensure that they were not proliferating during the 2-hour period of light-induced clustering. Epithelial gaps were almost absent in these non-proliferative stages (Figure S3B), which suggests that cell division challenges cell attachment in the follicular epithelium. Consistent with this, we observed frequent local loss of apical contacts next to dividing cells in *aPKC*^*as4*^ neuroepithelia upon aPKC inhibition (Figure S3C), despite the absence of large tissue rupture. To address how cell division contributed to loss of tissue integrity, we imaged epithelial gaps forming in the follicular epithelium stained with a membrane marker. Strikingly, the majority of epithelial gaps initiated as ruptures between cells undergoing cytokinesis and their neighbors (Figures 5A and 5B; Movie S6)

**Figure 5.**
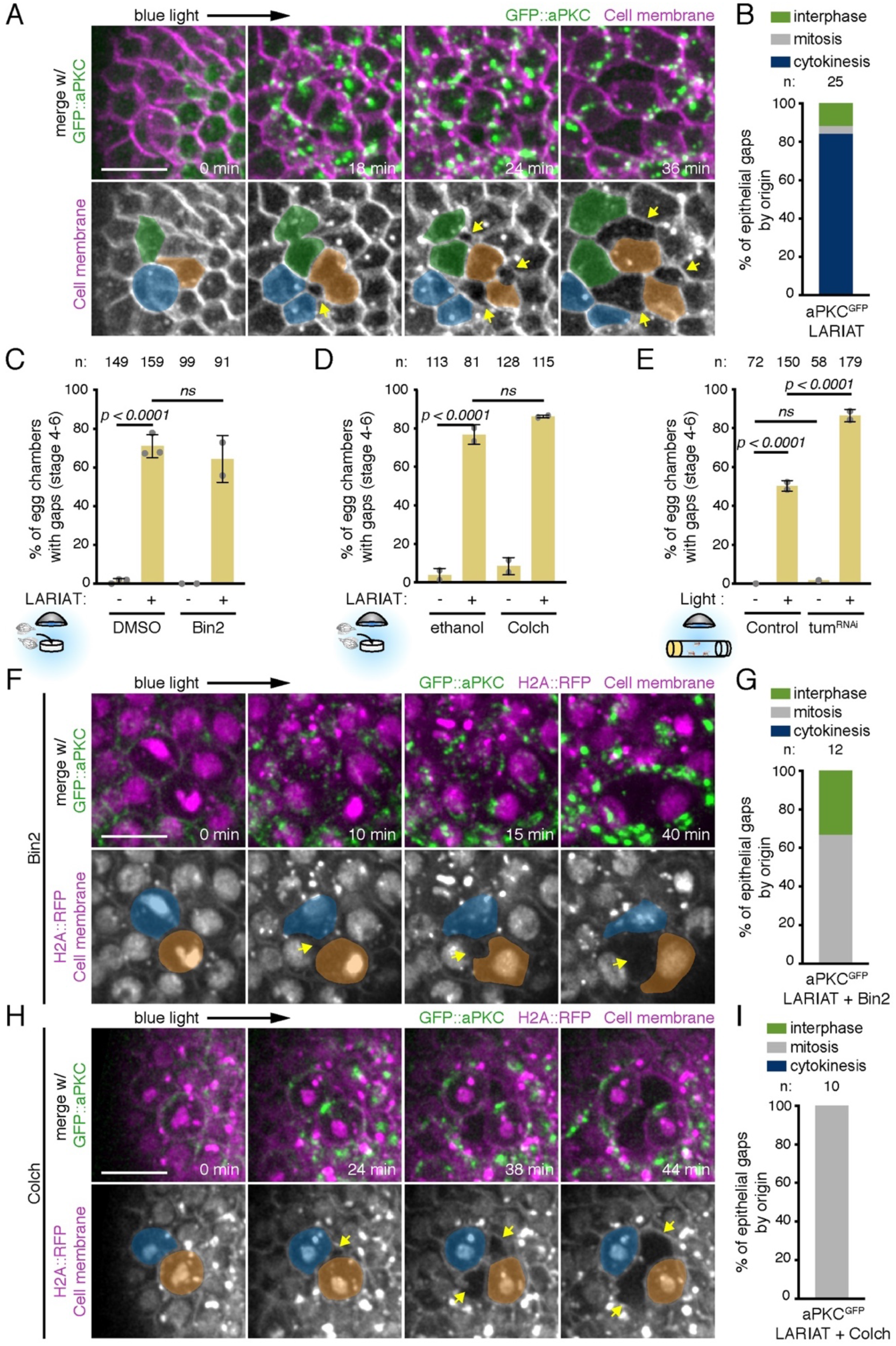
Cell division challenges tissue cohesion upon aPKC inactivation. **(A)** Time-lapse images of an egg chamber (surface view) expressing LARIAT, GFP::aPKC and stained with membrane marker. Imaging with 488nm laser clustered aPKC from min 0 onwards. Epithelial gaps (arrow) form adjacent to dividing cells (colored). **(B)** Quantification of epithelial gap origin according to cell division stage of neighbouring cells. n = number of gaps scored. **(C**,**D**) Gap frequency in the presence and absence of LARIAT in egg chambers treated with the indicated drug: **(C)** Binucleine-2 (Bin2), to inhibit AurB, or **(D)** Colchicine (Colch), to depolymerize microtubules, before light exposure for 2 hours *ex vivo*. **(E)** Frequency of epithelial gaps scored in control and Tum RNAi egg chambers from flies expressing GFP::aPKC LARIAT and exposed (+) or not (-) to blue light for 2 hours. Graphs in **(C-E)** show mean ± SD; grey data points represent independent experiments; n = number of egg chambers scored; Fisher’s exact test was used (ns, not significant). **(F-I)** Time-lapse images of follicular epithelium (surface views) expressing LARIAT, GFP::aPKC and H2A::RFP and stained with membrane marker. **(F)** Bin-2 or **(H)** Colch were added at least 15 min prior to clustering from min 0 onwards. Epithelial gaps (arrows) form adjacent to dividing cells (colored) despite **(F)** cytokinesis failure (chromatin (H2A::RFP) decondenses without chromosome separation) or **(H)** mitotic arrest (condensed chromatin throughout the movie). **(G**,**I)** Epithelial gap origin according to cell division stage of neighbouring cells. n = number of gaps scored. Scale bars: 10 µm. See also Figure S3.

The force produced by cytokinetic ring constriction could promote epithelial rupture in the context of aPKC downregulation by pulling on neighboring cells undergoing apical constriction. To test whether cytokinesis was necessary for epithelial cells to detach from each other, we blocked cytokinesis either by disrupting contractile ring assembly with the Aurora B inhibitor Binucleine 2 (Bin2) [52], or by blocking cells in prometaphase with the microtubule-depolymerizing drug Colchicine (Colch) (Figure S3D). We also depleted Tumbleweed/RacGAP50C (Tum; Figure S3E), a component of the centralspindlin complex that controls contractile ring assembly [53]. However, neither of these treatments prevented the formation of epithelial gaps upon aPKC clustering (Figures 5C-E). In addition, live imaging of egg chambers treated with Colch or Bin2 showed that optogenetic clustering of aPKC still led to recurrent tissue rupture next to mitotic cells even though they did not undergo cytokinesis (Figures 5G and 5H). Altogether, we conclude that while tissue rupture upon aPKC perturbation is commonly observed next to dividing cells, it does not require cytokinesis. Thus, other aspects of cell division likely provide an additional challenge to epithelial upon increased apical constriction.

### Dividing cells are stretched by hypercontractile neighbors after aPKC inactivation

To understand why epithelial gaps form between dividing and neighboring cells, we further characterized the impact of aPKC on MyoII distribution in mitotic and non-mitotic cells. During interphase, MyoII is enriched at the junctions and at the medioapical surface of follicle cells, where it drives pulses of apical constriction [54]. Live imaging showed GFP::aPKC accumulated at the apical intercellular contacts and displayed a smaller dynamic medioapical pool that accompanied the cycles of medioapical MyoII accumulation in interphasic follicle cells (Figures 6A and 6B; Movie S7). During mitosis, aPKC extends along the lateral cortex, a spatial redistribution that is also remininescent of MyoII (Figure 6A) [38]. To evaluate the effect of aPKC inactivation on distinct MyoII pools, we imaged Sqh::mKate2 in apical sections of epithelial cells. MyoII rapidly accumulated at the medioapical level upon optogenetic or chemical aPKC inhibition in interphasic follicle cells (orange arrows; Figures 6C and 6D; Movie S8). In contrast, in mitotic cells, aPKC inactivation did not affect the normal reduction of medioapical myosin (yellow arrows) nor the mitotic redistribution of myosin along the lateral cortex (blue arrow; Figures 6C and 6D). This result suggests that aPKC is required to antagonize apicomedial actomyosin specifically during interphase.

**Figure 6.**
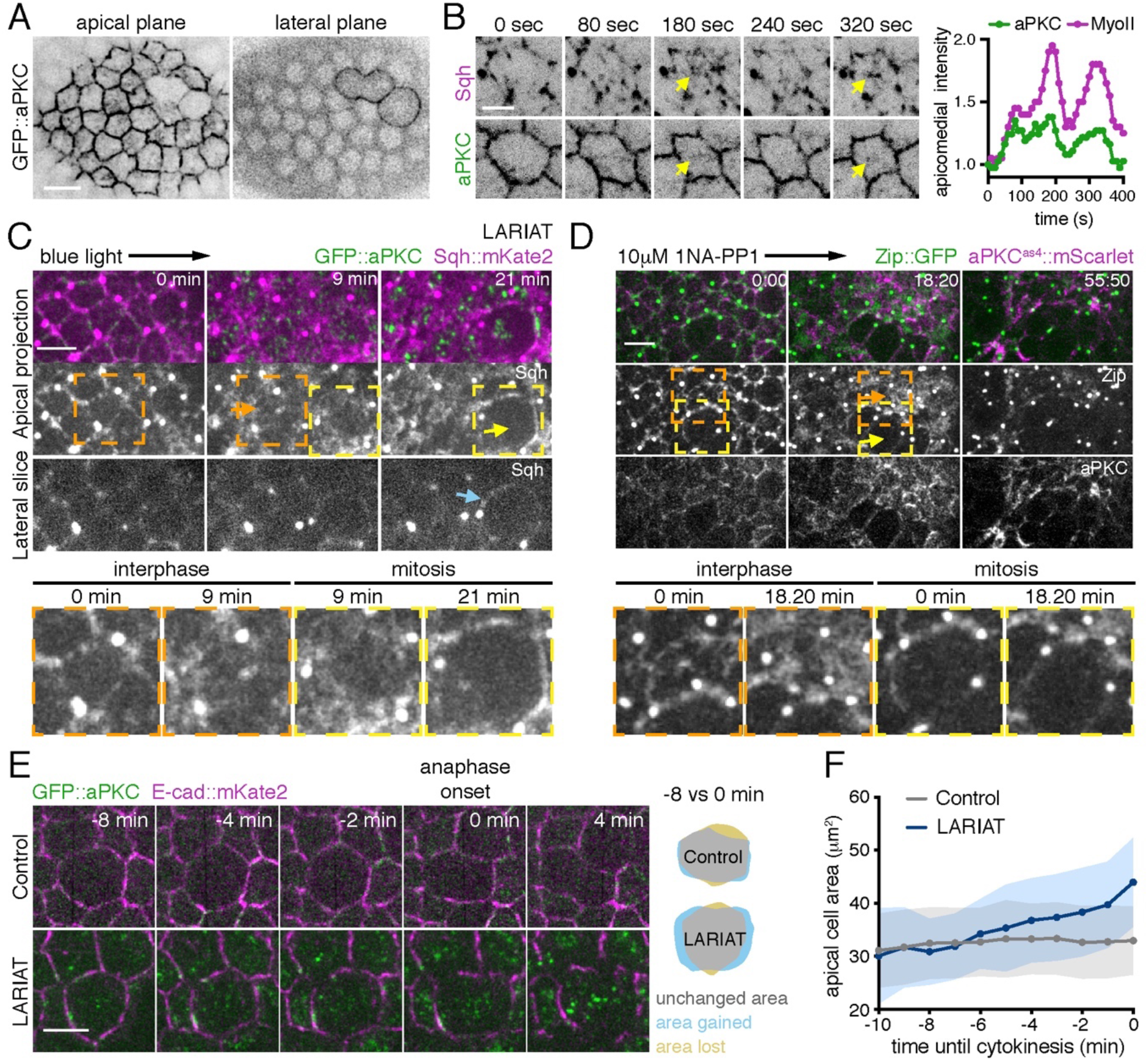
aPKC inactivation leads to excessive pulling forces on dividing cells. **(A)** GFP::aPKC in the follicular epithelium at the apical surface (left) and at the lateral cortex (right). **(B)** Live-imaging of GFP::aPKC and Sqh::mKate2 at the apical surface shows dynamic accumulation at the apicomedial region (arrows). Normalized, background subtracted mean pixel intensity of GFP::aPKC and Sqh::mKate2 measured in the medioapical region using a circular 2.5µm diameter ROI (right). **(C)** Live imaging of GFP::aPKC LARIAT in the follicular epithelium with Sqh::mKate2. Imaging with the 488 nm laser triggered aPKC clustering from min 0. After GFP::aPKC clustering, Sqh::mKate2 accumulates at the apicomedial region in interphase cells (orange arrow), but not in mitotic cells (yellow arrows), where aPKC clustering does not affect redistribution to the lateral cortex (blue arrow). **(D)** Time-lapse images of *aPKC*^*as4*^*::mScarlet* (magenta), Zip::YFP (green) follicle cells treated with 10 µM 1NA-PP1 at timepoint 0. Transient apicomedial accumulation of Zip::YFP (orange arrow) is not observed in mitotic cells (yellow arrow). **(C**,**D)** Insets show individual interphase and mitotic cells. **(E)** Live imaging of GFP::aPKC and E-cad::mKate2 in the follicular epithelium (apical projection). aPKC clustering was triggered up to 5 min before mitotic entry. Cells enter anaphase at min 0. **(F)** Apical surface area in dividing cells was measured at the junction level until anaphase onset (n = 27 cells from 6 control egg chambers and 12 cells from 8 LARIAT egg chambers. Graphs show mean ± SD. Scale bars: 5 µm.

We hypothesised that increased apical constriction in neighboring non-mitotic cells could produce excessive pulling on dividing cells, which could be unable to sustain this force due to the decrease in apical myosin. To evaluate this hypothesis, we monitored the effect of optogenetic aPKC inactivation on the apical surface area of mitotic cells with E-cad::mKate2. In contrast to interphasic cells (Figure 4A), mitotic cells did not contract, but rather expanded their apical domain upon mitotic entry and later detached from constricting neighboring cells (Movie S9). Quantification of the apical area in cells that were mitotic in the initial period of light exposure showed that clustering increased the expansion of the apical domain during mitosis (Figures 6E and 6F). Hence, excessive apical contractility in non-mitotic cells induces stretching of dividing cells and promotes tissue disruption.

### Increase in apical contractility at the tissue-level induces epithelial gaps

To address whether rupture of the follicular epithelium was produced by a global or local increase in apical contractility, we analysed proliferative epithelial tissue with clonal expression of UAS-LARIAT. In contrast to egg chambers where the whole tissue expressed UAS-LARIAT, there was no rupture in mosaic egg chambers containing cells insensitive to light (Figures 7A and 7B), neither if LARIAT cells divided adjacent to wild-type cells (n = 16) nor within LARIAT clones (n = 28). This result suggests that the apical side of wild-type cells may stretch to accommodate the constriction of neighboring tissue and prevent tissue rupture. Accordingly, wild-type cells neighboring UAS-LARIAT patches expanded their apical area during light exposure (Figure 7C).

**Figure 7.**
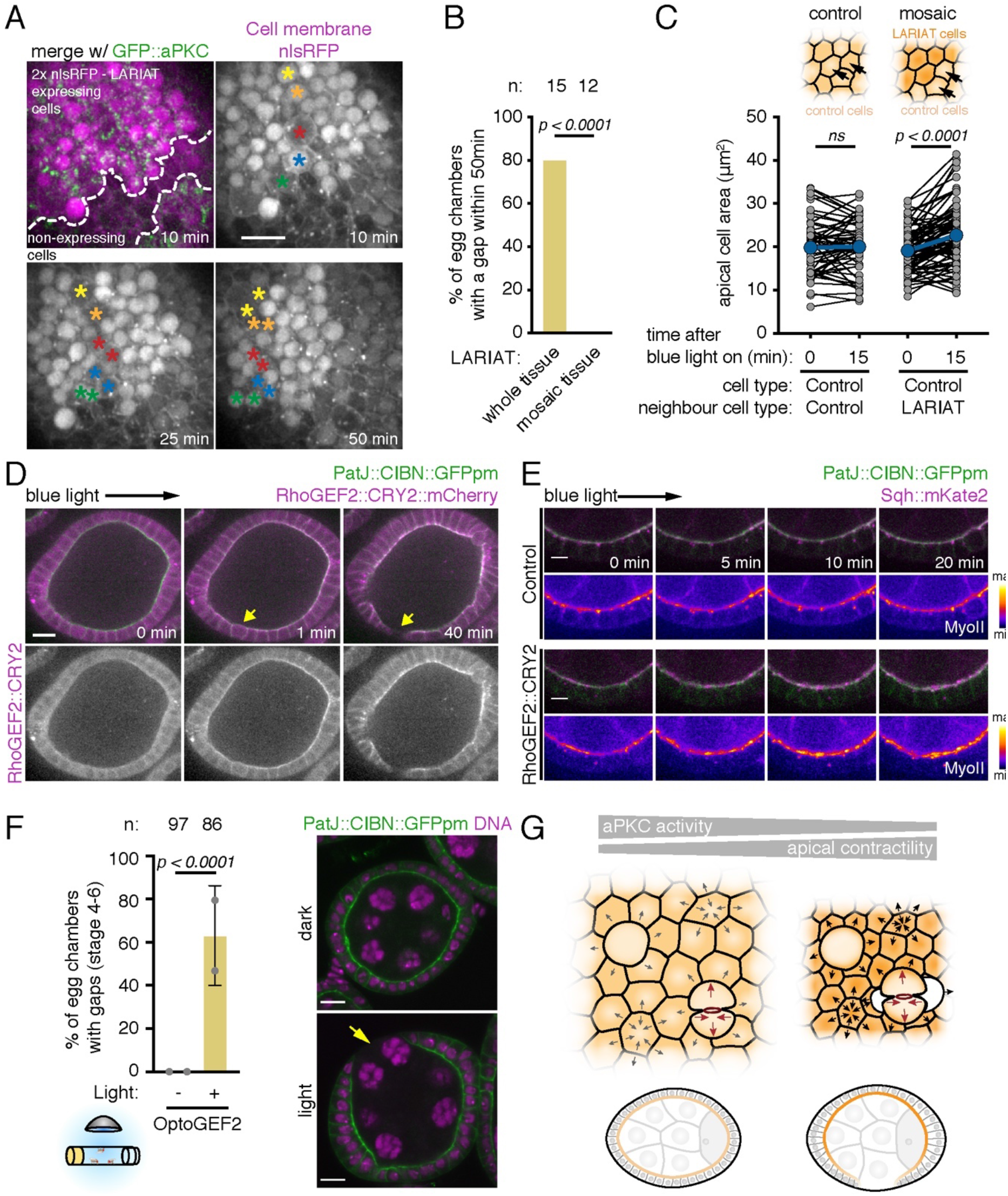
Global increase of apical contractility induces epithelial gaps. **(A)** Time-lapse images (surface of egg chamber) show epithelial rupture does not occur in GFP::aPKC follicular epithelia with mosaic LARIAT expression (marked by nls::RFP in magenta) despite multiple dividing cells (asterisks). **(B)** Frequency of gaps in tissue with mosaic LARIAT expression compared with whole tissue expression of LARIAT (data from Figure 5B re-analysed, Fisher’s test). **(C)** Variation off apical cell area shows that wild-type neighbour cells (adjacent to LARIAT-expressing clones, n = 82 cells, 13 egg chambers) expand within 15 min of aPKC clustering, unlike wild-type cells in a fully wild-type tissue (n = 60 cells, 2 egg chambers). Points represent individual cells (ANOVA for paired samples). **(D)** Live imaging of egg chamber expressing PatJ::CIBN::pmGFP and RhoGEF2::CRY2::mCherry in the follicular epithelium (midsagittal view). Imaging with 488 nm laser targeted RhoGEF2 to the apical domain from timepoint 0 onwards. Yellow arrow denotes gap formation in a region with a dividing cell. **(E)** Time-lapse images of the follicular epithelium expressing PatJ::CIBN::pmGFP, RhoGEF2::CRY2 and Sqh::mKate2 (midsagittal view). Pseudo-colored Sqh::mKate2 panel shows increase at the apical surface after RhoGEF2 apical recruitment. **(F)** Representative midsagittal images and epithelial gap (yellow arrow) quantification in egg chambers (stages 4 to 6) from flies expressing PatJ::CIBN::pmGFP and RhoGEF2::CRY2::mCherry exposed to blue light for 2 hours prior to fixation and staining for DNA. Control flies were left in the dark. Graphs show mean ± SD; grey data points represent independent experiments; n = total amount of egg chambers; Fisher’s exact test was used (ns, not significant); Scale bars: 5 µm. **(G)** Model depicts how downregulation of aPKC activity increases apical constriction in the non-mitotic cells of a polarized epithelium, generating excessive pulling forces on dividing cells that induce detachment and, ultimately, epithelial gaps.

To test if the global increase of apical contractility is sufficient to drive tissue rupture, we used an optogenetic tool to stimulate apical constriction by light-dependent recruitment of the RhoGTPase activator RhoGEF2 (RhoGEF2::CRY2::mCherry) to an apically enriched PatJ::CIBN::GFP::CAAX fusion [55]. Live imaging showed cytoplasmic RhoGEF2::CRY2::mCherry was quickly recruited to the apical domain after light exposure (Figure 7D). Apical recruitment of RhoGEF2::CRY2::mCherry induced apical MyoII accumulation and produced gaps next to dividing cells (Figures 7D and 7E). Moreover, *in vivo* exposure of flies expressing this optogenetic system to 2h of blue light reproduced the phenotype of aPKC LARIAT, leading to a high frequency of egg chambers with epithelial gaps (Figure 7F). Thus, increased apical contractility is sufficient to disrupt epithelial integrity in a proliferative epithelium, further supporting the idea that aPKC protects epithelial integrity through regulation of apical actomyosin.

## Discussion

Apical-basal polarity provides positional information at the cellular level that is essential for tissue architecture. However, it remains ill-defined how loss of polarity regulators leads to epithelial architecture defects. Even though genetic approaches have yielded substantial insight, the inherent temporal constraints preclude direct visualization of the underlying events. Here, we used fast aPKC perturbation approaches in *Drosophila* epithelia to shed light on how aPKC regulates epithelial architecture. We show that epithelial gaps form prior to loss of apical-basal polarity and within minutes of aPKC downregulation in the follicular epithelium. aPKC inactivation increases apical contractility in non-mitotic cells. This increase pulls dividing and neighbor cells apart and ruptures the epithelium. Thus, we propose that aPKC downregulates apical contractility in polarized epithelia to prevent the build-up of excessive forces that compromise epithelial integrity (Figure 7G).

Rapid protein perturbation approaches are necessary to define the primary cellular cause of phenotypes in tissues. Here, we have developed a strategy to quickly inactivate aPKC in epithelial and neural stem cells by employing optogenetic clustering in the abdomen of living flies or *ex vivo* in intact organs. By complementing optogenetic clustering with the ability to adjust aPKC activity with chemical-genetics, we show that, immediately after clustering, aPKC is only partially inactive. More importantly, this allowed us to show that decreasing aPKC activity initially increases apical contractility and leads to the formation of gaps in the follicular epithelium. Strikingly, partial inactivation did not disrupt apical-basal polarity immediately, revealing that this is not the direct cause of epithelial gaps. This recapitulates the phenotype of hypomorphic *aPKC* alleles that produce gaps but do not disrupt apical-basal polarity [11, 45]. Our results therefore indicate that a high threshold of aPKC inactivation is required to disrupt apical-basal polarity, which suggests that polarized epithelia can withstand fluctuations in aPKC activity. In turn, the higher sensitivity of the apical actomyosin cytoskeleton likely enables aPKC-dependent regulation of contractility without compromising apical-basal polarity.

aPKC is essential for apical-basal polarity, which provides spatial cues that orient the organization of the apical actomyosin network. However, aPKC has also been reported to antagonize the apical actomyosin network during morphogenesis in *Drosophila* and mammalian embryos [35, 56, 57]. We now propose that aPKC downregulates apical contractility to balance forces within proliferating epithelia to maintain epithelial integrity. Interestingly, whereas inhibition of aPKC induces accumulation of myosin at the apicomedial surface in follicle cells, myosin increase is more pronounced at junctions in neuroepithelial tissue. Consequently, our work highlights a primary role for aPKC as a negative regulator of apical constriction whose function can be locally regulated for different morphogenetic and homeostatic purposes. The molecular nature of this function has yet to be uncovered. Phosphorylation of ROCK by aPKC induces its cortical dissociation to downregulate junctional contractility in mammalian cells [26], but aPKC does not regulate equivalent sites in *Drosophila* Rok [58]. Alternatively, aPKC may target other actomyosin regulators or function through other apical polarity proteins implicated in the regulation of apical contractility, such as Crumbs and Lulu2/Yurt [34, 59-63].

Our findings suggest that physical constraints also define the phenotypic outcome of apical constriction. Our analysis shows that epithelial gaps form almost exclusively at the dorsal/ventral regions of egg chambers, where tension at the apical domain has been reported to be higher [54, 64]. The follicular epithelium is physically constrained at the basal side by a stiff basement membrane [65] and at the apical side by the growing germline, which may keep the epithelium stretched [66, 67]. Thus, egg chamber organization likely opposes the shape changes necessary to accommodate global apical constriction, leading to an increase in tension and rupture upon aPKC inactivation. In contrast, neuroepithelial cells constrict their apical area freely, which should release tension. Accordingly, while partial aPKC inactivation consistently led to large epithelial gaps in the follicular epithelium, in the neuroepithelium only minor perturbations at apical cell contacts were detected. Hence, on top of possible differences in the local response at the junctional or medioapical level, our findings stress the importance of physical boundaries, tissue geometry and mechanical context on the outcome of increased apical contractility.

We also provide direct evidence that large epithelial rupture can arise by intercellular detachment during cell division, which provides a weak spot primed for disruption upon increased mechanical stress. This is consistent with previous observations that reinforcement of junctional attachment to the cytoskeleton prevents detachment during cell division in the *Drosophila* embryonic epithelium and mammalian cell culture [68, 69]. Dividing cells do not generate gaps upon aPKC inactivation in a mosaic tissue, showing that a direct effect in mitotic cells is not responsible for gap formation by itself. Then, why are mitotic cells prone to separate from hypercontractile surrounding tissue? Mitotic cells downregulate apicomedial actomyosin and revert apical constriction, which makes them more susceptible to extrinsic forces [70, 71]. We show that increased pulling forces exerted by the constricting non-mitotic tissue indeed expand the apical surface of mitotic cells upon aPKC clustering. These forces could amplify outward pulling forces at the poles of dividing cells [72], and spatially oppose pulling forces by the contractile ring on cell adhesion during cytokinesis. We observed that ruptures generally occur next to the equatorial region during cytokinesis, which is consistent with opposing forces overcoming cell adhesion in this region (Figure 7G). Furthermore, local remodelling of cell adhesion during mitosis [39, 73, 74] and cytokinesis [38, 75-78] may favour detachment next to dividing cells.

Different cellular events have to be integrated at the tissue level to drive concerted shape changes during morphogenesis. Apical constriction is frequently used to bend or fold epithelia during development [79]. Cell division actively contributes to tissue morphogenesis by controlling tissue material properties [74, 80-82] and driving shape change [83-85] or cellular rearrangements [86]. However, the cell-intrinsic mitotic remodelling of the cytoskeleton can disrupt morphogenetic processes that require apical constriction [70, 87-89]. Our results now show that forces produced by apical constriction challenge cohesion at the dividing-neighboring cell interface, and thereby disrupt epithelial integrity. Hence, this study shows the importance of a strict control over apical constriction in proliferative tissues, so as to enable growth and morphogenesis without compromising epithelial integrity.

## Acknowledgments

We thank Daniel St Johnston, Juergen A. Knoblich, Stefano de Renzis, Xiaobo Wang, Yohanns Bellaiche and the Bloomington *Drosophila* Stock Center for fly stocks. We also thank Yohanns Bellaiche, Ivo Telley and Romain Levayer for insightful comments on the manuscript. This work was funded by National Funds through FCT—Fundação para a Ciência e a Tecnologia, I.P., under the projects PTDC/BEX-BCM/0432/2014 and PTDC/BIA-CEL/1511/2021. E.M. is funded by “FCT Scientific Employment Stimulus - Individual Call” program (CEECIND/00622/2017). M.O. and A.M.C. were supported by PhD fellowships from FCT through the GABBA PhD program. We also acknowledge the support of the i3S Scientific Platform ALM, member of the national infrastructure PPBI - Portuguese Platform of Bioimaging (PPBI-POCI-01-0145-FEDER-022122). Work in JJ’s lab is supported by the 100031/Z/12/Z and 100031/Z/12/A from Wellcome and the Royal Society. We would like to thank the Dundee Imaging Facility for excellent support.

## Author contributions

E.M. and M.O. conceptualized the study and wrote the original manuscript draft; Data acquisition: M.O., A.B., A.M.C. and E.M. performed all experiments in follicular epithelium apart of the experiment performed by N.L. in Fig. 6D; N.L performed experiments in larval neuroepithelium; P.G. performed neuroblast experiments. Data analysis and interpretation: M.O., A.B., A.M.C., N.L., P.G., C.H, J.J., and E.M.; Supervision: C.S., C.H., J.J. and E.M. Funding acquisition: C.H, J.J. and E.M.; All authors reviewed the manuscript.

## METHODS

### EXPERIMENTAL MODEL AND SUBJECT DETAILS

#### Drosophila melanogaster genetics and husbandry

We performed all experiments using *Drosophila melanogaster*. We raised fly lines on standard fly food (cornmeal/agar/molasses/yeast) at 18°C or 25°C with 60% humidity and 12h/12h dark light cycle, except when otherwise indicated in the method details section.

The following fly lines were used:

- Under regulation of the respective endogenous promoters: *Par-6::GFP* ([90], gift from Jürgen Knoblich, IMBA, Vienna), *H2A::RFP* ([91] and *Sqh::mKate2×3* – *Drosophila* non-muscle myosin II regulatory light chain tagged with three tandem *mKate2* and inserted in chromosomes II and III ([92], gift from Yohanns Bellaïche, Institut Curie, Paris);
- Tagged in the respective endogenous locus: *GFP::aPKC* ([43], gift from Daniel St Johnston, The Gurdon Institute, Cambridge), *Lgl::mCherry [22]* and *E-cad::GFP* [93], both gifts from Yang Hong, University of Pittsburgh), *E-cad::mKate2×3* and *E-cad::GFPx3* (tagged with three tandem *mKate2* or *GFP*, [92], gift from Yohanns Bellaïche, Institut Curie, Paris) and *Zip::YFP* – *Drosophila* non-muscle myosin II heavy chain [94];
- *UAS-LARIAT* inserted in chromosomes II and III: *CRY2* PHR domain fused to SNAPtag and a GFP nanobody (*V*_*H*_*H*) and N-terminal *CIB* domain (residues 1-170) fused to the *CaMKIIα* multimerization domain (MP), both expressed from a single construct with the help of a P2A self-cleaving peptide ([44], gift from Xiabo Wang, CBI, Université de Toulouse);
- For optogenetic RhoGEF2 recruitment to the apical membrane: *PatJ::CIBN::pmGFP*, for UAS-driven expression of *PatJ* fused to N-terminal *CIB* domain (residues 1-170) and membrane targeted *GFP* fused to human *Kras4B* CAAX [55], *RhoGEF2::CRY2* and RhoGEF2::CRY2::mCherry, for UAS-driven expression of tagged and untagged catalytic DHPH domain of *RhoGEF2* fused to *CRY2* PHR [95], gifts from Stefano de Renzis, EMBL, Heidelberg);
- To drive Gal4-UAS mediated construct expression: *GR1-Gal4*, an enhancer trap line where Gal4 is under regulation of unknown regulatory sequences that drive Gal4 expression in the follicular epithelium ([96] and *tj-Gal4*, an enhancer trap line where Gal4 is under regulation of *traffic jam* regulatory sequences (DGGR_104055); *pnt-Gal4* (VDRC_VT212057, discarded)
- *Gal80*^*ts*^, temperature-sensitive *Gal80* under regulation of the αTub84B promotor (BDSC_7018);
- nlsRFP hs-Flp FRT19A (BDSC_31418) and hs-Flp Gal80 FRT19A (BDSC_5133), for FLP/FRT-mediated generation of Gal80 clones.
- *aPKC*^*k06403*^, an *aPKC* null allele obtained by insertional mutagenesis of a P-element construct [6];
- *aPKC*^*as4*^ is an ATP analog-sensitive aPKC allele (I342A and T405A mutations introduced in the endogenous locus through CRISPR/Cas9 [48];
- aPKC::mScarlet^as4^ was made by scar-less (inDroso co-CRISPR approach) CrispR gene editing. The mScarlet-I sequence, preceded and followed by short two amino acid (VAL GLY) linkers, was inserted into the genome of the *apkc*^*as4*^ line using the AATGGATCCTCCGGTGGCGGTGG guide RNA. The insert position was the same as previously published for GFP [97]: the mScarlet-I amino acid sequence, including the ATG and framed by the linkers, was inserted after amino acid 228 of aPKC-PA.
- UAS-mCherry (BDSC_35787), UAS-Tum RNAi (BDSC_28982) and phosphomimetic and nonphosphorylatable Sqh - UAS-Sqh^E20E21^ (BDSC_64411) and UAS-Sqh^A20A21^ (BDSC_64114);

Fly genotypes for each experiment can be found in **Table S1**. For optogenetic experiments where flies were exposed to blue light, female offspring of the same cross with the same genotype were randomly assigned to experimental groups (dark vs light). For each independent *ex vivo* experiment with drug treatment of egg chambers, we dissected ovaries from all flies of the same genotype, mixed them together, separated their ovarioles and then randomly distributed them by the experimental groups. For live imaging of egg chambers, we imaged 2-3 egg chambers per fly.

#### Optogenetic experiments in the follicular epithelium

To inactivate apical polarity with optogenetics, we combined *aPKC* tagged with GFP in its endogenous locus [43] with *UAS-LARIAT* [44]. We used *tj-Gal4* or *GR1-Gal4* to drive *UAS* constructs expression in the follicular epithelium. To minimize premature UAS construct expression, crosses were kept at 18°C. 1-3 days after eclosion, adult offspring were transferred to 29°C to drive expression of UAS constructs (1 day at 29°C for all UAS constructs, except for optogenetic RhoGEF2 membrane recruitment (Figure 7D-F, which was induced for 2-3 days at 29°C). To avoid unintended optogenetic system activation by light, we kept fly vials inside cardboard boxes or wrapped in aluminum foil and handled them in a dark room under a 593nm LED light source (SuperBrightLEDs) from this point forward. For co-expression of LARIAT with other UAS constructs, we used temperature-sensitive *Gal80*^*ts*^ to fully suppress premature Gal4-UAS driven transcription prior to temperature shift to 29°C. To express LARIAT in clones, we generated *Gal80* clones through FLP/FRT-mediated recombination [98]. These crosses were kept at 18°C, protected from light and heat shocked at 37°C for 2 hours 3-5 times. LARIAT-expressing cells were marked by the presence of 2 nlsRFP copies, while wild-type cells had either 1 or no nlsRFP copies. Alternatively, in Figure S2A, LARIAT-expressing cells were identified by the presence of GFP::aPKC clusters.

For *in vivo* optogenetic experiments, flies were exposed continuously to blue light for the indicated periods of time (in Table S1 and figure legends) by placing vials at approximately 8 cm from a 472nm LED bulb (SuperBrightLEDs) at room temperature. Afterwards, we dissected their ovaries and fixed them. For each independent experiment, control flies from each genotype were kept in the dark and dissected in a dark room to avoid triggering CIBN-CRY2 interaction. To control for potential blue light toxicity, we also exposed flies without optogenetic constructs to blue light using the same setup (data included in Figure 1).

For *ex vivo* optogenetic experiments, ovaries were dissected in a dark room in *ex vivo* culture medium (Schneider’s medium (Sigma-Aldrich) supplemented with 10% FBS (fetal bovine serum, heat inactivated; Thermo Fisher) and 200 μg/mL insulin (Sigma-Aldrich)). Afterwards, the dissected ovaries were transferred to new *ex vivo* culture medium and the ovarioles were partially separated by pipetting up and down gently. The separated ovarioles were exposed to blue light for 2 hours in 24-well-plates using the same setup as for whole flies and then they were fixed and stained to evaluate epithelial architecture. When indicated in the figures and figure legends, specific drugs (or DMSO or etanol for control samples) were added 20 minutes before exposure to blue light: Colchicine (Sigma-Aldrich; 30 μM; prepared in ethanol) to depolymerize microtubules and block cells in mitosis; Binucleine-2 (Sigma-Aldrich; 40 μM; prepared in DMSO) to inhibit Aurora B; and Latrunculin A to disrupt the actin cytoskeleton (Sigma Aldrich; 5 μg/mL; prepared in DMSO). To confirm that Binucleine-2 blocked cytokinesis and Colchicine blocked cells in mitosis in the follicular epithelium, ovarioles were treated with these drugs for 30 minutes and then fixed (without exposing them to blue light). For live imaging, ovaries were dissected in a dark room and CIBN-CRY2 interaction was only triggered with the 488 nm laser used for GFP-tagged protein imaging.

#### Fixation and staining of egg chambers

To evaluate epithelial architecture, *Drosophila* ovaries were dissected in Schneider’s medium (Sigma-Aldrich) supplemented with 10% FBS (fetal bovine serum, heat inactivated; Thermo Fisher), washed once with PBT (PBS + 0.05% Tween 20 (Sigma-Aldrich)) and fixed in 4% paraformaldehyde solution (prepared in PBS with 0.2% Tween 20 (Sigma-Aldrich)) for 20 min. For F-actin staining, Phalloidin-FITC (Sigma-Aldrich) or Phalloidin-TRITC (Sigma-Aldrich) was added to the fixative solution at 1 μg/mL and incubation time was increased to 30 min. After washing three times for 10 min with PBT, samples were mounted with Vectashield with DAPI (Vector Laboratories). To evaluate mitotic progression (for Figure S3D), egg chambers were stained for Histone H3 phosphorylated Ser10 and actin. After fixation and washing, samples were blocked for 2 hours at room temperature with 10% FBS (prepared in PBS + 0.2% Tween 20) and incubated overnight at 4°C with rabbit anti-phospho-Histone H3 (pH3) Ser10 (1:2000; Upstate Biotechnology) diluted in PBT + 1% FBS. Afterwards, the samples were washed four times with PBT + 1% FBS for 30 minutes and incubated for two hours at room temperature with Alexa Fluor 568-conjugated goat anti-rabbit (Invitrogen; 1:300) diluted in PBT + 0.1% FBS. After washing two times with PBT for ten minutes, samples were incubated for 30 minutes with Phalloidin-TRITC diluted in PBT, washed three times for ten minutes and mounted with Vectashield with DAPI (Vector Laboratories).

#### Optogenetic experiment and immunofluorescence in neuroblasts

We used *pnt-Gal4* to drive *UAS-LARIAT* expression in type II neuroblasts. Following a 12h egg-laying period, control and LARIAT embryos were kept in the dark until wandering L3 larvae (wL3) stage. wL3 of both conditions were exposed to light for 1h. Brains were then dissected in PBS 1x, fixed in 4% paraformaldehyde for 20 min at room temperature and washed three times with PBST (0.1% Triton X-100 in 1x PBS). Brains were blocked with 1% normal goat serum in 0.1% PBST for at least 20 min at room temperature and incubated overnight at 4°C with rabbit anti-Miranda (1:2000, [99], gift from Juergen A. Knoblich) and mouse monoclonal anti-phospho-Histone H3 (pH3) Ser10 (1:1000, Cell Signalling, 9706), diluted in blocking solution. Afterwards, brains were washed three times, blocked for 20 min and incubated for 2h at room temperature with secondary antibodies Alexa Fluor 647-conjugated goat anti-mouse and Alexa Fluor 568-conjugated goat anti-rabbit (Invitrogen), used at 1:1000. Finally, brains were mounted in Aqua Polymount (Polysciences, Inc).

#### Imaging of fixed tissue

Images of fixed *Drosophila* egg chambers were collected with a 1.1 NA/40x water or 1.30 NA/63x glycerine objectives on an inverted laser scanning confocal microscope Leica TCS SP5 II (Leica Microsystems) or 1.30 NA/63x glycerol objective on an inverted laser scanning confocal microscope Leica SP8 (Leica Microsystems). To score epithelial defects and evaluate mitotic progression, images for egg chamber staging were collected with a 10x objective on a Zeiss Axio Imager Z1 microscope (Carl Zeiss, Germany) or a Zeiss Axio Imager Z1 Apotome microscope (Carl Zeiss, Germany). To evaluate epithelial architecture defects (epithelial gaps and/or multilayering), midsagittal cross-sections of egg chambers were inspected with a 20x or 40x Oil objective. To evaluate mitotic progression, images from the follicular epithelium at the surface of egg chambers were acquired with a 40x Oil objective on a Zeiss Axio Imager Z1 microscope (Carl Zeiss, Germany). Images from *Drosophila* larvae brains were acquired with a Zeiss LSM880 confocal microscope (Zeiss) using a LD LCI Plan-Apochromat 40x/1.2 Imm Corr DIC M27 water objective.

#### Live imaging

For live imaging of *Drosophila* egg chambers, individual ovarioles were dissected in *ex vivo* culture medium (Schneider’s medium (Sigma-Aldrich) supplemented with 10% FBS (fetal bovine serum, heat inactivated; Thermo Fisher) and 200 μg/uL insulin (Sigma-Aldrich)) and the enveloping muscle removed as previously described. Ovarioles were transferred to new culture medium and imaged on uncoated coverslips or glass bottom dishes (MatTek; No 1.5; P35G-1.5-7-C) with an Andor XD Revolution Spinning Disk Confocal system equipped with two solid state lasers – 488nm and 561nm -, an iXonEM+ DU-897 EMCCD camera and a Yokogawa CSU-22 unit built on an inverted Olympus IX81 microscope with a PLAPON 60x/NA 1.42 or a UPLSAPO 100x/NA 1.40 objective using iQ software (Andor). On average 2 egg chambers were imaged per fly. When indicated in the figures, to mark the cell membrane, ovarioles were stained with CellMask Orange Plasma membrane Stain (ThermoFisher; C10045; diluted 1:10 000 in culture medium) for 15 minutes and washed twice with *ex vivo* culture medium before imaging. Live imaging was performed at 25°C. When indicated in the figures and figure legends, Colchicine (Sigma-Aldrich; 30 μM; prepared in etanol), Binucleine-2 (Sigma-Aldrich; 40 μM; prepared in DMSO) or Latrunculin A (Sigma Aldrich; 5 μg/mL; prepared in DMSO) were added at least 15 minutes before imaging. Midsagittal egg chamber cross-sections were used to image the follicular epithelium along the apical-basal axis and z-stacks at the surface of the egg chamber to cross-section the follicular epithelium along the apical-basal axis.

For live imaging of larval brain neuroepithelia, brains from L3 larvae were dissected in Schneider’s medium supplemented with glucose (1 mg/ml) and insulin (0.2 mg/ml) and transferred to a 10 μl drop of the same medium supplemented with Fibrinogen (0.2 mg/ml) on a 25 mm glass-bottom dish. Brains were oriented on their side and the Fibrinogen was clotted using thrombin (100 U/ml, Sigma-Aldrich, T7513). After 3 min, 190 μl Schneider’s medium supplemented with glucose and insulin was pipetted on top of the clot. The neuroepithelia were imaged for 15 minutes on a Zeiss 710 Spinning Disk microscope using a 63x Plan-Apochromat 1.4 NA objective. 200 μl Schneider’s medium supplemented with glucose, insulin and 1NA-PP1 (20 μM) was then added for a final concentration of 10 μM 1NA-PP1, after which imaging was resumed.

#### Protein extracts and Western blot

To confirm endogenous and GFP::aPKC levels in the different genotypes used for optogenetic aPKC inactivation (Figure S1A), we prepared protein extracts from *Drosophila* ovaries (at least 15 flies per genotype) dissected in a dark room. Dissected *Drosophila* ovaries were transferred to lysis buffer (150mM KCl, 75mM HEPES pH 7.5, 1.5 mM EGTA, 1.5mM MgCl_2_, 15% glycerol, 0.1% NP-40, 1x protease inhibitors cocktail (Roche) and 1x phosphatase inhibitors cocktail 3 (Sigma-Aldrich)), frozen in liquid nitrogen, thawed and then disrupted through sonication. We clarified lysates through two consecutive centrifugations at 14000 rpm for 10 min at 4°C. Protein concentration was determined with NanoDrop 1000 Spectrophotometer (Thermo Fisher). Samples were then resolved through SDS-PAGE and transferred to a nitrocellulose membrane using the iBlot Dry Blotting System (Invitrogen) for Western blotting. Protein transfer was confirmed by Ponceau staining (0.25% Ponceau S in 40% methanol and 15% acetic acid). The membranes were blocked for two hours at room temperature with 5% dry milk prepared in PBT and incubated overnight at 4°C with the primary antibodies (rabbit anti-aPKC 1:2000 (c-20, Santa Cruz Biotechnology) and mouse anti-α-Tubulin 1:10 000 (DM1A, Santa Cruz Biotechnology)) diluted in PBT + 1% dry milk. After washing three times for 10 min with PBT, membranes were incubated with the secondary antibodies anti-mouse and anti-rabbit conjugated with horseradish peroxidase diluted in PBT + 1% dry milk for one hour at room temperature. After washing again three times for 10 min with PBT, blots were developed with ECL Chemiluminescent Detection System (Amersham) according to the manufacturer’s instructions and revealed with a ChemiDoc XRS+ (BioRad).

#### *aPKC*^*as4*^ allele inactivation

For epithelial defect analysis in the follicular epithelium, *Drosophila* ovaries from aPKC^as4^ flies (prepared as previously described in the optogenetic experiments section) were cultured *ex vivo* for 2 hours in the presence of the ATP analog 1NA-PP1 (Calbiochem; prepared in DMSO; at the concentrations indicated in Figure 2 and the respective figure legend) before fixation. DMSO was added to control samples. For live imaging of egg chambers and larval neuroepithelium, 1NA-PP1 (at the concentration indicated in figure legends) or DMSO was added to the culture medium at the indicated timing.

### QUANTIFICATION AND STATISTICAL ANALYSIS

Image processing and quantifications were done with FIJI [100]. Statistical analysis and graphs were done in GraphPad Prism 8 (GraphPad Software Inc., La Jolla, CA, USA), except when otherwise indicated.

#### Epithelial defects analysis

To evaluate epithelial architecture, we scored the amount of egg chambers at specific developmental stages with one or more epithelial gaps, one or more multilayering events or both in midsagittal cross-sections. As egg chambers develop, they grow in size. Thus, we determined the developmental stage of egg chambers by measuring their area in midsagittal cross-sections, as a proxy for size. To define the area intervals corresponding to each developmental stage, we staged control egg chambers from *GFP::aPKC* flies according to phenotypic characteristics, as in Jia *et al*., [101], and correlated their stage with their size. We scored epithelial defects (epithelial gaps and/or multilayering) and their position (anterior, posterior, dorsal-ventral) by inspecting midsagittal cross-sections of egg chambers: for LARIAT and aPKC^as4^ experiments, egg chambers were stained with DAPI (DNA) and Phalloidin (F-actin); for optogenetic RhoGEF2 membrane recruitment, egg chambers were stained with DAPI (DNA) and PatJ-CIBN-pmGFP and RhoGEF2-CRY2-mCherry fluorescence was used. For the initial analysis of aPKC inactivation with LARIAT (Figure 1C), results from 3 independent experiments (≥ 8 flies per condition per experiment) were summed up in a single contingency table and the graph shows the relative amount of egg chambers (stages 3 to 8) with each type of defect found. For statistical analysis, epithelial gaps and/or multilayering were grouped in a single defect category and Fisher’s exact test with Bonferroni correction for multiple comparison was used. For other experiments, graphs show mean percentage of egg chambers with the indicated type of defect ± standard deviation (SD). The percentages of defective egg chambers obtained for each independent experiment for each condition (≥ 8 flies per condition per independent experiment) are represented as individual data points in the graphs. The total amount of egg chambers scored in each analysis is indicated in the respective graph as n. To ensure consistent LARIAT expression levels, only proliferative stages 4 to 6 were included in analyses (except in Figures 1 and S3B). To test for statistical significance, we built contingency tables comparing the sum of egg chambers from all replicates with one or more epithelial gap vs no gap and used Fisher’s exact test, with Bonferroni correction for multiple comparisons when necessary. To compare the frequency of epithelial gap and multilayering formation at the anterior, posterior and dorsal-ventral regions, we analyzed how many of the epithelial gaps and multilayering events detected upon aPKC clustering in proliferative egg chambers were present at these different regions irrespective of how long the samples had been exposed to blue light (data in Fig. 2C).

#### Lgl asymmetry in the follicular epithelium

The ratio of apical over lateral Lgl fluorescence intensity was used to analyse the asymmetric distribution of Lgl along the apical-basal axis in the follicular epithelium after aPKC clustering. For each egg chamber, the lateral and apical cortex of all epithelial cells was manually segmented, average fluorescence intensity was extracted and corrected for average Lgl intensity in the cytoplasm. Parts of the epithelium presenting with multilayering were excluded from the analysis (at least 10 cells per egg chamber were included in the analysis). Each point in Figure 1E represents the apical/lateral Lgl ratio for an individual egg chamber.

#### Miranda asymmetry in neuroblasts

The distribution of Miranda along the membrane of dividing NBs (pH3 positive) was analysed. To extract Miranda intensity profile, that is, the intensity of Miranda (I_M_) along the length of the cell membrane (L), we proceeded as in (Rodriguez et al., 2017) with minor changes: we used a 30-pixel wide stripe to delineate the membrane; profile extraction was initiated in the basal membrane section, so that L<50% correspond to basal Miranda intensity values. In the end, Miranda intensity (I_M_) was plotted as a function of percentage of membrane length (L). To obtain the asymmetry index (ASI) for Miranda, Basal (B) and Apical (A) intensities were calculated as the area under the Miranda intensity plot (I_M_) for the basal (L ≤50%) and apical (L>50%) sections of the membrane. Absolute ASI values were calculated as in (Hannaford et al., 2019) with the following formula:

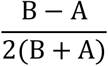

ASI values were then normalized relative to control mean so that, an ASI of 1 represents normal asymmetry and lower values (∼0) indicate loss of asymmetry. As *pnt*-GAL4 only drives LARIAT expression in type II neuroblasts, only this subtype was considered in all calculations. Statistical analysis was performed using Prism 6 (GraphPad Software Inc., La Jolla, CA, USA), mean ± SD are depicted and individual ASI values represented (normalized to control mean). Statistical significance of the difference of means was calculated using unpaired *t-*test and considered significant when *p* < 0.05.

#### Apical myosin II in the follicular epithelium

To quantify apical accumulation of myosin II after aPKC inactivation, we measured Sqh::mKate2×3 fluorescence intensity at the apical domain of follicular epithelial cells in single plane midsagittal cross-sections of egg chambers acquired during live imaging. For each egg chamber, 4 regions of interest (ROIs), each one encompassing the apical domain of at least 4 cells in a different region (dorsal, ventral, anterior or posterior), were manually defined and tracked through time. Mean apical fluorescence intensity (AFI) for each timepoint was extracted from raw movie datasets, corrected for mean cytoplasm fluorescence intensity (CFI; average value measured for each timepoint in 3 follicular epithelial cells) and normalized to the corrected fluorescence intensity before aPKC inactivation (AFI_initial_-CFI_initial_) as follows:

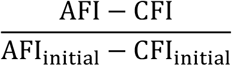

For aPKC LARIAT experiments, AFI_initial_-CFI_initial_ corresponds to the AFI-CFI value at min 0 (when optogenetic clustering is triggered). For aPKC^as4^ experiments AFI_initial_-CFI_initial_ was obtained by averaging AFI-CFI for the 5 frames before aPKC inactivation with 1NA-PP1 at min 0. Whenever, an epithelial gap started forming at a particular region, quantification at that same region was stopped.

#### Egg chamber circularity

To assess egg chamber deformation after aPKC inactivation, we measured egg chamber circularity in single plane midsagittal cross-sections of Sqh::mKate2×3 egg chambers acquired during live imaging. The apical surface of the follicular epithelium was manually segmented and circularity (4π (area/perimeter^2^)) was measured. Egg chamber circularity was only quantified while no epithelial defect appeared.

#### Apical surface area and myosin II in the neuroepithelium

To measure apical surface contraction in the neuroepithelium upon aPKC^as4^ inactivation with 1NA-PP1, the ROI edges were manually tracked as red dots using aPKC^as^-mScarlet signal to detect apical cell edges. These dots were then connected using a steerable filter for line detection. The resulting shape was then filled and dilated (blue mask in Figure 3G) to approximate the area to measure. Apical area was normalized to its initial value at min 0. To measure myosin II intensity at the apicomedial region and apical junctions after aPKC^as4^ inactivation with 1NA-PP1, apical junctions were segmented using aPKC^as^-mScarlet signal. aPKC signal to noise ratio was increased by a steerable filter detecting lines and a junctional mask (as the one shown in Figure 3J) was generated by thresholding. The mask for the apicomedial region was generated by inverting and contracting the junctional mask. Myosin II intensity was obtained by extracting average Zip::YFP intensity with these masks and normalizing to junctional intensity at min -2 before aPKC inactivation.

#### Apical area in interphase and mitotic cells in the follicular epithelium

To evaluate apical constriction in interphase cells and pulling forces on mitotic cells, we measured epithelial cell area in cross-sections at the junctional level of the follicular epithelium acquired during live imaging. For each egg chamber, we quantified the mean apical cell area (average of at least 3 interphase cells manually segmented using E-cad::mKate2 per egg chamber). Surface area was normalized to the initial mean value, obtained by averaging the corresponding cell area for the 3 frames before aPKC clustering (from min -2 to 0). A similar procedure was used to segment cells that entered mitosis up to 5 minutes after aPKC clustering was initiated. Anaphase onset was defined as the first frame of cell elongation (determined through E-cad::mKate2 signal at the lateral cortex). Mitotic entry was defined as the first frame of visible mitotic rounding in a lateral cortex cross-section.

To evaluate apical constriction in clones of LARIAT-expressing cells, we measured the epithelial cell area at the apical surface of the follicular epithelium acquired during live imaging. For each clone, we quantified the mean apical cell area (average of at least 6 cells per clone manually segmented in 4D stacks using GFP::aPKC and a plasma membrane marker). A similar procedure was used to measure the apical area of individual wild-type cells adjacent to LARIAT-expressing cells (Figure 7C).

#### Epithelial gap analysis live

To evaluate whether and where gaps formed in the follicular epithelium, 4D stacks of surface cross-sections from egg chambers stained with membrane marker were analyzed. Gaps were inspected to verify if they span the whole length of the apical-basal axis and were only included in the analysis when all neighbour cells were in sight, so as to be able to determine whether any of them were in mitosis. The number of independent gaps detected in the 13 control, 8 Binucleine-2-treated and 6 Colchicine-treated egg chambers is indicated as n in Figures 5B, 5G and 5I, respectively.

#### Mitotic progression

To confirm the effect of Binucleine-2 and Colchicine, we analyzed mitotic progression in control and drug treated egg chambers. Mitotic cells were identified through positive staining with pH3 (number of cells counted indicated as n in Figure S3D and E). DAPI staining was used to verify whether sister chromatids had separated and group cells into early mitosis (prophase, prometaphse, metaphase) and late mitosis (anaphase, telophase) or cytokinesis. DNA. Actin staining was used to verify whether cells had elongated, confirming anaphase onset, and whether they had assembled a cytokinetic ring.

#### Image preparation

Representative images were processed and prepared using FIJI. Representative midsagittal images from egg chambers are from a single optical section or 2-5 plane maximum projection. Surface images from egg chambers are maximum projections of all optical sections covering the epithelial domain of interest. When necessary, movies were registered with the FIJI plugin *StackReg* (EPFL; Biomedical Imaging Group), to correct for whole egg chamber movement during live imaging; and a Gaussian Blur or Gaussian Blur 3D filter was applied to remove random noise.

