## Supplementarydata for "Apical constriction induces tissue rupture in a proliferative epithelium"

**Figure S1: Optogenetic clustering of GFP::aPKC disrupts Miranda asymmetry in neural stem cells. Related to Figure 1**

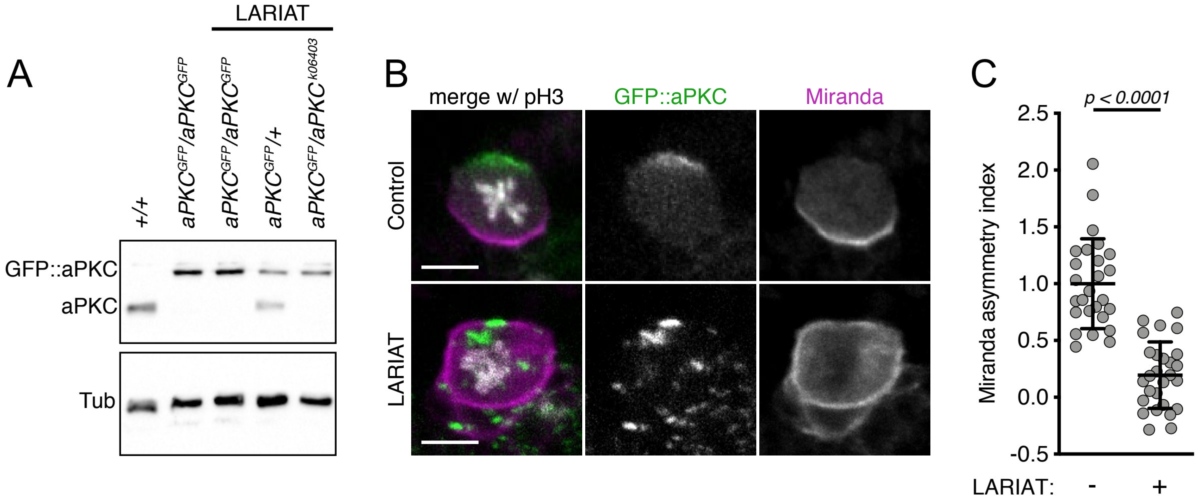


**(A)** Western blot shows protein level for untagged and GFP::aPKC in ovaries for the indicated genotypes used in Figure 1C. $\alpha$-Tubulin was used as loading control. **(B)** Representative images of control and LARIAT neuroblasts stained for Miranda (magenta) and pH3 to label mitotic cells (white). Exposure to blue light for 1 hour clustered GFP::aPKC and prevented release of Miranda from the apical cortex. **(C)** Miranda asymmetry index along the cell cortex in control (n = 25 cells) and LARIAT neuroblasts (n = 27 cells) was normalized to control mean value. Graph shows mean ± SD values (p<0.0001, t-test). Scale bars: 5 µm.

**Figure S2: Acute aPKC inactivation does not disrupt the junctional accumulation of E-cad and upregulates apical myosin. Related to Figure 3.**

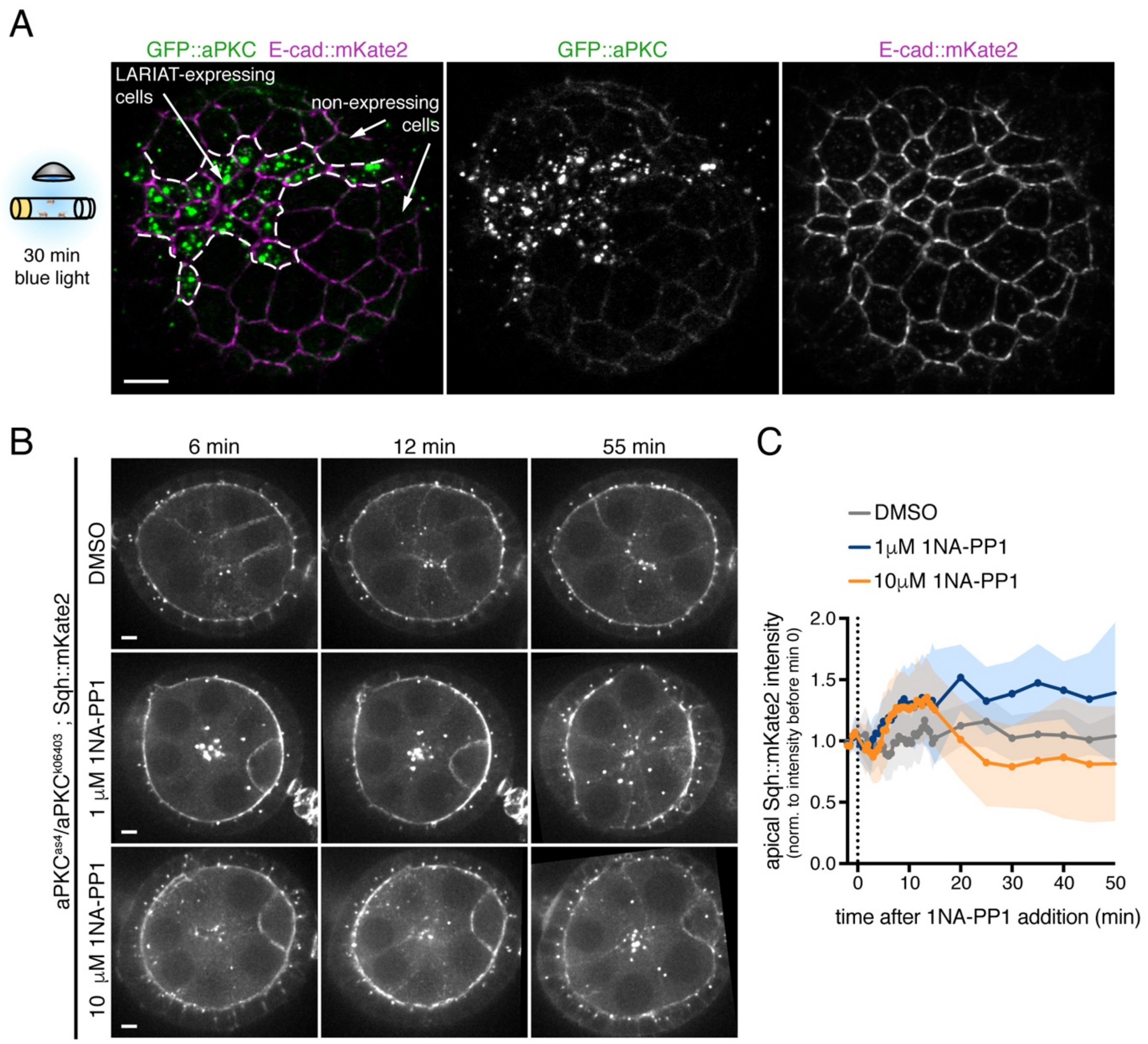


**(A)** Confocal Z-stack projection showing an egg chamber (surface view) that expresses endogenously tagged E-cad::mKate2 and contains mosaic clones of UAS-LARIAT cells that cluster GFP::aPKC (green). Exposure of flies to blue light for 30 min reduces the apical area of LARIAT expressing follicle cells but does not disrupt the junctional accumulation of E-cad (magenta). **(B)** Time-lapse midsagittal images of *aPKC*$^{as4}$*/aPKC*$^{k06403}$ egg chambers expressing Sqh::mKate2. The indicated concentrations of 1NA-PP1 were added at timepoint 0. **(C)** Sqh intensity at the apical surface was corrected for cytoplasm intensity and normalized to average intensity prior to aPKC inhibition (n ≥ 32 cells from ≥ 2 egg chambers per condition). Scale bars: 5 µm.

**Figure S3: Formation of epithelial gaps during cell division upon aPKC inactivation. Related to Figure 5.**

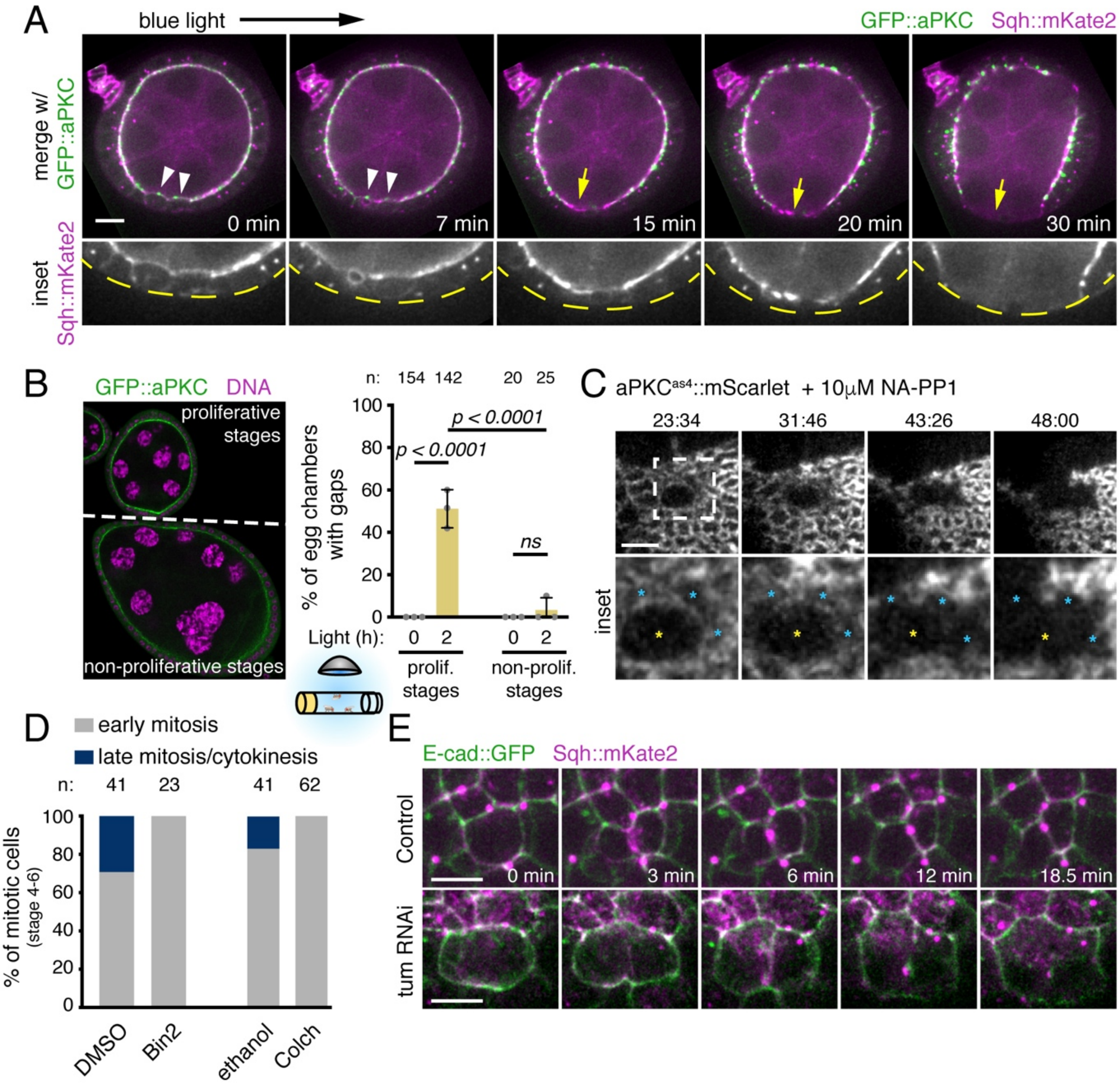


**(A)** Time-lapse images of egg chambers (midsagittal view) expressing LARIAT, GFP::aPKC (green) and Sqh::mKate2 (magenta). Imaging with 488nm laser clustered aPKC from min 0 onwards. Epithelial gap (yellow arrow) forms where cells divided (white arrowheads); dashed line delineates egg chamber in Sqh inset. **(B)** Quantification of epithelial gaps in proliferative (stages 4-6) and non-proliferative (analysis restricted to stage 8 egg chambers to ensure the sample was non-proliferative during the 2h of light exposure) GFP::aPKC homozygous egg chambers expressing LARIAT and exposed to light *in vivo* for 0 or 2h before ovary fixation. Graphs show mean ± SD ($p<0.0001$, Fisher's test); grey data points represent percentage from independent experiments; n = number of egg chambers scored. **(C)** Time-lapse images of *aPKC$^{as4}$::mScarlet* shows dividing (yellow asterisk) and non-dividing neighbour cells (blue asterisk) in the neuroepithelium treated with 10 μM 1NA-PP1 at timepoint 0. Loss of apical contacts (marked by aPKC) after division occurred in 8/16 divisions in 9 epithelia. **(D)** Quantification of the frequency of mitotic cells in early vs late mitosis (anaphase onwards) in egg chambers treated with Binucleine-2 (Bin2), Colchicine (Colch) or respective controls confirms that both drugs disrupt cytokinesis initiation. Mitotic cells were scored in egg chambers stained for phospho Ser10 on Histone H3, actin and DNA. n = number of mitotic cells scored**.** **(E)** Live imaging of cell division in follicle cells expressing Sqh::mKate2 and E-cad::GFP in control and Tum RNAi egg chambers. Tum-depleted cells start to constrict but fail cytokinesis (cytokinesis onset at min 0). Scale bars = (A) 10 μm (C,E) 5 μm.

**Table S1. Genotypes and light exposure details** (related to STAR methods)**.**

*Drosophila* genotype for each figure panel. When applicable, optogenetic experiment details are indicated: whether optogenetic system was activated by exposing intact flies or dissected ovaries cultured *ex vivo* to blue light and how long samples were exposed to blue light. For live imaging experiments, optogenetic system was only activated after image acquisition started.

| **Genotype** | | **Optogenetic system activation details** |
|---|---|---|
| **Figure 1** | | |
| B | ; *GFP::aPKC*; *GR1>Gal4/UAS>LARIAT*<br>; *GFP::aPKC/+*; *GR1>Gal4/UAS>LARIAT* | Flies – 2 days |
| D | ; *tj>Gal4, GFP::aPKC/GFP::aPKC*; (Control)<br>; *tj>Gal4, GFP::aPKC/GFP::aPKC*; *UAS>LARIAT/+*<br>; *tj>Gal4, GFP::aPKC/+*; *UAS>LARIAT/+*<br>; *tj>Gal4, GFP::aPKC/*$aPKC^{k06403}$; *UAS>LARIAT/+* | Flies – 1 day |
| E | *; GFP::aPKC, Lgl::mCherry/GFP::aPKC*; *UAS>LARIAT/+*<br>; *tj>Gal4, GFP::aPKC/GFP::aPKC, Lgl::mCherry*; *UAS>LARIAT/+* | Flies – 1 day |
| **Figure 2** | | |
| A | ; *GFP::aPKC*; *GR1>Gal4/UAS>LARIAT* | Flies – 0, 2, 4, 6 or 12 hours |
| B,C | ; *tj>Gal4, GFP::aPKC/GFP::aPKC*; *UAS>LARIAT/+* | Flies – 0, 2, 4, 6 or 12 hours |
| D | ; *tj>Gal4, GFP::aPKC/GFP::aPKC*; *H2A::RFP/UAS>LARIAT* | Live imaging |
| E, F | ; $aPKC^{as4}$; | N/A |
| **Figure 3** | | |
| A | ; *tj>Gal4, GFP::aPKC/Lgl::mCherry, GFP::aPKC*; *UAS>LARIAT/+*<br>; *tj>Gal4 GFP::aPKC/GFP::aPKC, E-cad::mKate2x3*; *UAS>LARIAT/+* | Live imaging |
| B-D | ; *tj>Gal4, GFP::aPKC/GFP::aPKC*; *Sqh::mKate2x3/UAS>LARIAT* | Live imaging |
| E,F | *ubi::nlsRFP, hsFlp, FRT19A/hsFlp, tub>Gal80, FRT19A*; *tj>Gal4 GFP::aPKC/GFP::aPKC, UAS>LARIAT*; +<br>LARIAT expressing cells are marked by bright RFP signal (two nlsRFP copies) and GFP::aPKC clustering. | Live imaging |
| G, H | ; $aPKC^{as4}$*::mScarlet*; | N/A |
| I, J | ; $aPKC^{as4}$*::mScarlet, Zip::YFP/*$aPKC^{as4}$*::mScarlet* | N/A |

| | | |
|---|---|---|
| **Figure 4** | | |
| A,B | *; GFP::aPKC, E-cad::mKate2x3/GFP::aPKC; UAS>LARIAT/+*<br>*; tj>Gal4, GFP::aPKC/GFP::aPKC, E-cad::mKate2x3; UAS>LARIAT/+* | Live imaging |
| C,D | *; tj>Gal4, GFP::aPKC/GFP::aPKC; +*<br>*; tj>Gal4, GFP::aPKC/GFP::aPKC; UAS>LARIAT/+* | Ovaries *ex vivo* – 2 hours |
| E | *; tj>Gal4, GFP::aPKC/GFP::aPKC, UAS>LARIAT; Gal80*$^{ts}$*/UAS>mCherry*<br>*; tj>Gal4, GFP::aPKC/GFP::aPKC, UAS>LARIAT; Gal80*$^{ts}$ */UAS>Sqh*$^{E20E21}$<br>*; tj>Gal4, GFP::aPKC/GFP::aPKC, UAS>LARIAT; Gal80*$^{ts}$ */UAS>Sqh*$^{A20A21}$ | Flies – 2 hours |
| **Figure 5** | | |
| A | *; GFP::aPKC; GR1>Gal4/UAS>LARIAT* | Live imaging |
| B, F-I | *; tj>Gal4, GFP::aPKC/GFP::aPKC; H2A::RFP/UAS>LARIAT* | Live imaging |
| C, D | *; tj>Gal4* GFP::aPKC/GFP::aPKC;<br>*; tj>Gal4* GFP::aPKC/GFP::aPKC; UAS>LARIAT/+ | Ovaries *ex vivo* – 2 hours |
| E | *; tj>Gal4, GFP::aPKC/GFP::aPKC, UAS>LARIAT; Gal80*$^{ts}$*/UAS>mCherry*<br>*; tj>Gal4, GFP::aPKC/GFP::aPKC, UAS>LARIAT; Gal80*$^{ts}$*/UAS>Tum RNAi* | Flies – 2 hours |
| **Figure 6** | | |
| A | *; tj>Gal4, GFP::aPKC/GFP::aPKC;* | N/A |
| B | *; tj>Gal4, GFP::aPKC/GFP::aPKC; Sqh::mKate2x3/+* | N/A |
| C | *; tj>Gal4, GFP::aPKC/GFP::aPKC; Sqh::mKate2x3/UAS>LARIAT* | Live imaging |
| D | *; aPKC*$^{as4}$*::mScarlet, Zip::YFP/aPKC*$^{as4}$*::mScarlet* | N/A |
| E-F | *; GFP::aPKC, E-cad::mKate2x3/GFP::aPKC; UAS>LARIAT/+*<br>*; tj>Gal4, GFP::aPKC/GFP::aPKC, E-cad::mKate2x3; UAS>LARIAT/+* | Live imaging |
| **Figure 7** | | |
| A, C | *ubi::nlsRFP, hsFlp, FRT19A/hsFlp, tub>Gal80, FRT19A; tj>Gal4 GFP::aPKC/GFP::aPKC, UAS>LARIAT; +*<br>LARIAT expressing cells are marked by bright RFP signal (two nlsRFP copies) and GFP::aPKC clustering. | Live imaging |
| B | *; tj>Gal4, GFP::aPKC/GFP::aPKC; H2A::RFP/UAS>LARIAT* (LARIAT expression in whole tissue)<br>*ubi::nlsRFP, hsFlp, FRT19A/hsFlp, tub>Gal80, FRT19A; tj>Gal4 GFP::aPKC/GFP::aPKC, UAS>LARIAT; +* (mosaic LARIAT expression) | Live imaging |
| D | *; tj>Gal4/UAS>PatJ::CIBN::pmGFP; UAS>RhoGEF2::CRY2::mCherry/+* | Live imaging |
| E | *; tj>Gal4/UAS>PatJ::CIBN::pmGFP; UAS>RhoGEF2::CRY2/Sqh::mKate2x3* | Live imaging |
| F | *; tj>Gal4/UAS>PatJ::CIBN::pmGFP; UAS>RhoGEF2::CRY2::mCherry/+* | Flies – 2 hours |
| **Supplementary Figure S1** | | |

| | | |
|---|---|---|
| A | $w^{1118}$;;<br>; *tj>Gal4, GFP::aPKC/GFP::aPKC*;<br>; *tj>Gal4, GFP::aPKC/GFP::aPKC*; *UAS>LARIAT/+*<br>; *tj>Gal4, GFP::aPKC/+*; *UAS>LARIAT/+*<br>; *tj>Gal4, GFP::aPKC/aPKC*$^{k06403}$; *UAS>LARIAT/+* | dark |
| B, C | ; *GFP::aPKC/GFP::aPKC*; *pnt>GAL4/MKRS*<br>; *GFP::aPKC, UAS>LARIAT/GFP::aPKC*; *pnt>GAL4/MKRS* | Larvae – 1 hour |
| **Supplementary Figure S2** | | |
| A | *FRT19A/hsFlp, tub>Gal80, FRT19A*; *tj>Gal4, GFP::aPKC, E-cad::mKate2/GFP::aPKC, UAS>LARIAT*; +<br>LARIAT expressing cells are marked by GFP::aPKC clustering. | Flies 30 min |
| B,C | ; *aPKC*$^{as4}$ /aPKC$^{k06403}$, *E-cad::GFPx3*; *sqh::mKate2x3/+* | N/A |
| **Supplementary Figure S3** | | |
| A | ; *tj>Gal4 GFP::aPKC/GFP::aPKC*; *Sqh::mKate2x3/UAS>LARIAT* | Live imaging |
| B | ; *tj>Gal4, GFP::aPKC/GFP::aPKC*; *UAS>LARIAT/+* | Flies – 0, 2 hours |
| C | ; *aPKC*$^{as4}$*::mScarlet*; | N/A |
| D | ; *tj>Gal4, GFP::aPKC/GFP::aPKC*; *UAS>LARIAT/+* | ovaries *ex vivo*, dark |
| E | ; *tj>Gal4, E-cad::GFP, Sqh::mKate2x3/+*;<br>; *tj>Gal4, E-cad::GFP, Sqh::mKate2x3/+*; *Gal80*$^{ts}$*/UAS>Tum RNAi* | Live imaging |

**Supplementary Movie Legends:**

**Movie S1, related to Figure 2. aPKC inactivation leads to fast tissue disorganization.** Live imaging of egg chamber (midsagittal view) expressing GFP::aPKC (green), LARIAT, H2A::RFP (magenta), and stained with membrane marker (magenta). Imaging with 488nm laser triggered GFP::aPKC clustering from timepoint 0 onwards. Epithelial gaps appear at the dorsal and ventral side and multilayering at the posterior side. Right panel shows H2A::RFP and membrane marker.

**Movie S2, related to Figure 3. aPKC inactivation leads to rapid apical myosin accumulation and cell deformation.** Time-lapse movies shows that Sqh::mKate2x3 (magenta and as separate channel) increases at the apical membrane during the initial period of GFP::aPKC (green) clustering. Imaging with 488 nm laser triggered GFP::aPKC clustering from timepoint 0 onwards.

**Movie S3, related to Figure 3. aPKC inactivation induces cell-autonomous apical constriction.** Live imaging of GFP::aPKC (green) follicular epithelial cells (surface view) with mosaic expression of LARIAT (marked by the presence of two copies of nlsRFP (bright magenta) and aPKC clustering). Imaging with 488 nm laser triggered GFP::aPKC clustering from timepoint 0 onwards. Right panel shows GFP::aPKC.

**Movie S4, related to Figure 3. aPKC inactivation leads to apical constriction in the *Drosophila* larval neuroepithelium of the optic lobe.** Live imaging of the neuroepithelium expressing $aPKC^{as4}$::mScarlet. $aPKC^{as4}$ was inactivated with 10 $\mu$M 1NA-PP1 at timepoint 0.

**Movie S5, related to Figure 5. Epithelial gaps are generated next to dividing cells upon aPKC clustering (midsagittal view).** Live imaging of egg chamber (midsagittal view) expressing GFP::aPKC (green), LARIAT, Sqh::mKate2x3 (magenta). Imaging with 488nm laser triggered GFP::aPKC clustering from timepoint 0 onwards. Right panel shows Sqh::mKate2x3. Arrrowheads in the first frame point to dividing cells.

**Movie S6, related to Figure 5. Tissue rupture by intercellular detachment next to dividing cells (surface view).** Live imaging of egg chamber (surface view) expressing GFP::aPKC (green), LARIAT and stained with membrane marker (magenta). Imaging with 488nm laser triggered GFP::aPKC clustering from timepoint 0 onwards. Epithelial gap appears between dividing cells and their neighbours. Right panel shows membrane marker.

**Movie S7, related to Figure 6. aPKC has a dynamic medioapical pool that accompanies cycles of myosin accumulation.** Live imaging of GFP::aPKC (right panel) and MyoII(sqh)::mKate2 (left panel) in the follicular epithelium (surface view).

**Movie S8, related to Figure 6. Myosin accumulates at the medioapical level upon optogenetic inactivation of aPKC.** Live imaging of GFP::aPKC (green; right panel) and Sqh::mKate2 (magenta; middle panel) in the follicular epithelium (surface view). Imaging with 488nm laser triggered GFP::aPKC clustering in the presence of LARIAT from timepoint 0 onwards.

**Movie S9, related to Figure 6. Mitotic cells expand and are pulled apart from their neighbours by apical constriction in interphasic cells .** Live imaging of a GFP::aPKC (green; right panel) and E-cad::mKate2 (magenta) in the follicular epithelium (surface view). Imaging with the 488nm laser triggered GFP::aPKC clustering from timepoint 0 onwards. Unlike interphase cells (blue asterisks) constrict, mitotic cells (yellow asterisks) still expand after aPKC clustering.

**Movie S10, related to Figure 7. Mosaic aPKC inactivation does not produce epithelial rupture.** Live imaging of a GFP::aPKC (green) follicular epithelium with mosaic expression of LARIAT (marked by the presence of two copies of nlsRFP (bright magenta) and aPKC clustering) and stained for a membrane marker (magenta). Imaging with 488 nm laser triggered GFP::aPKC clustering from timepoint 0 onwards.

**Movie S11, related to Figure 7. Apical recruitment of RhoGEF2 induces tissue rupture next to dividing cells.** Imaging with 488nm laser triggered RhoGEF2::CRY2::mCherry recruitment to apical PatJ::CIBN::pmGFP (green).
